# Brain-wide genetic mapping identifies the indusium griseum as a prenatal and shared target of pharmacologically-unrelated psychostimulants

**DOI:** 10.1101/759423

**Authors:** Janos Fuzik, Sabah Rehman, Fatima Girach, Andras G. Miklosi, Solomiia Korchynska, Gloria Arque, Roman A. Romanov, János Hanics, Ludwig Wagner, Konstantinos Meletis, Yuchio Yanagawa, Gabor G. Kovacs, Alán Alpár, Tomas G.M. Hökfelt, Tibor Harkany

## Abstract

Psychostimulant use is an ever-increasing socioeconomic burden, including a dramatic rise during pregnancy. Nevertheless, brain-wide effects of psychostimulant exposure are incompletely understood. Here, we performed Fos-CreER^T2^-based activity mapping, correlated for pregnant mouse dams and their fetuses with amphetamine, nicotine and caffeine applied acutely during mid-gestation. While light-sheet microscopy-assisted intact tissue imaging revealed drug- and age-specific neuronal activation, the indusium griseum (IG) appeared indiscriminately affected. By using GAD67^gfp/+^ mice we subdivided the IG into a dorsolateral domain populated by GABA interneurons and a ventromedial segment containing glutamatergic neurons, many showing drug-induced activation and sequentially expressing *Pou3f3/Brn1* and secretagogin (Scgn) during differentiation. We then combined Patch-seq and circuit mapping to show that the ventromedial IG is a quasi-continuum of glutamatergic neurons (IG-Vglut1^+^) reminiscent to dentate granule cells in both rodents and humans, whose dendrites emanate perpendicularly towards, while their axons course parallel with the superior longitudinal fissure. IG-Vglut1^+^ neurons receive Vglut1^+^ and Vglut2^+^ excitatory afferents that topologically segregate along their somatodendritic axis. In turn, their efferents terminate in the olfactory bulb, thus being integral to a multi-synaptic circuit that could feed information antiparallel to the olfactory-cortical pathway. In IG-Vglut1^+^ neurons, prenatal psychostimulant exposure delayed the onset of Scgn expression. Genetic ablation of Scgn was then found to sensitize adult mice towards methamphetamine-induced epilepsy, suggesting a role for this Ca^2+^-binding protein in scaling IG-Vglut1^+^ neuronal excitability. Overall, our study identifies brain-wide targets of the most common psychostimulants, among which Scgn^+^/Vglut1^+^ neurons of the IG link limbic and olfactory circuits.

**Significance statement:** Drug abuse during pregnancy is a significant socioeconomic problem. The use of psychostimulants is particularly common during pregnancy even though a risk to the developing fetus is significant. Here, we show that short-lived exposure to amphetamine, nicotine and caffeine during pregnancy induces neuronal activation in the fetal brain with the indusium griseum (IG), a brain area situated parallel to the central surface of the cortical hemispheres, becoming indiscriminately activated. By using mouse genetics, we find that psychostimulants preferentially target glutamatergic IG neurons, and delay their differentiation postnatally. Notably, the expressional onset of secretagogin, a Ca^2+^-sensor amenable for synaptic integration, is deregulated. This is significant because these neurons are integral to a multi-synaptic neuronal pathway that links limbic and olfactory circuits. As such, genetic deletion of secretagogin brings about heightened sensitivity to psychostimulants, manifesting as epileptiform discharges. Cumulatively, we describe a novel psychostimulant-sensitive neuronal subtype and its circuit arrangement whose developmental delay seems critical for behavioral abnormalities in offspring prenatally exposed to the most common psychostimulants.

## Introduction

Drug abuse is a major socioeconomic burden, which affects many age groups and both genders (1, 2). Psychostimulants are amongst the most widely used drugs, and besides illicit substances like amphetamine and its derivatives, include caffeine (coffee) and nicotine (tobacco). Molecular and structural determinants mediating the action of these drugs on neurons differ fundamentally: amphetamine is excitatory by increasing extracellular dopamine levels through inhibition (or even reversal) of dopamine transport for cellular re-uptake (1, 3, 4). Caffeine antagonism at the adenosine A2A receptor (5) (and less so at A1 and A3) is a catalyst for catecholamine (particularly dopamine) and acetylcholine release (6). Nicotine activates ionotropic nicotinic acetylcholine receptors to stimulate synaptic neurotransmission, particularly in the dopaminergic reward circuitry (7). Despite differences in their mechanism of action, a common denominator of psychostimulant action is their ability to prime the brain’s reward circuitry to trigger addiction (2). Therefore, any brain region that receives significant dopamine input is at risk upon excess exposure to psychoactive substances.

The consumption of illicit or legal drugs during pregnancy is a primary health concern. Besides counteracting medical conditions, such as *hyperemesis gravidarum* by cannabis and cannabinoids (8), the use of psychostimulants is reasoned by improving forgetfulness, mood, and stress. Nevertheless, the efficient cross-placental transfer of these structurally distinct compounds (9–11) prompts caution for their direct, indiscriminate and likely adverse effect, particularly if used repeatedly. Mechanistically, dopaminergic neurons and their long-range projections to the forebrain form during the first trimester of human brain development (12), thereby being poised to modulate the migration and neuritogenesis of telencephalic neurons over long periods (13, 14). Nevertheless, the brain-wide distribution of fetal neurons that are particularly sensitive to psychostimulants and differences between neural foci of fetal *vs*. maternal drug sensitivity remain ambiguous.

Here, we used immediate early gene-based mapping of transient cellular activity (15) to mark cell contingents (particularly neurons and endothelial cells) that were activated by amphetamine, nicotine or caffeine during the period when neurogenesis peaks in the mouse prosencephalon (that is, embryonic days (E) 14-16) (16, 17), and reconstructed their distribution by quantitative light-sheet microscopy (18). Besides finding critical differences in the regional density of genetically-tagged cell contingents, the medial preoptic area (MPN), arcuate nucleus (ARC) of the hypothalamus and the indusium griseum (IG), all receiving dense dopamine afferentation or containing dopaminergic neurons themselves (19), emerged as invariably affected fetal brain regions. This genetic model also revealed substantial differences in brain-wide cell activation patterns between dams and their fetuses.

We have then synthesized previous (20) and present findings on neuronal heterogeneity of the IG to show that glutamatergic neurons expressing the *Pou3f3/Brn1* transcription factor and many showing migratory morphology respond to amphetamine, caffeine and nicotine at the doses used. Once arriving to their final location in the ventromedial IG, an area dominated by excitatory neurons (20), postnatal neuronal differentiation is delayed by transient intrauterine psychostimulant exposure: we identified secretagogin (*Scgn*) as a psychostimulant-sensitive molecular mark for excitatory corpus callosum-associated neurons, which exhibit bipolar morphology in both the fetal and adult IG in mouse and humans. This is significant since the human IG was earlier claimed to be exclusively made up by glial cells (21). Next, we used *Patch-seq* to define cell type-specific marks for these regular-spiking (RS) *Scgn*^+^ neurons, such as *Vglut1* expression. By bi-directional connectivity mapping we also showed that *Scgn*^+^ IG*-Vglut1* neurons preferentially innervate GABA interneurons locally, while reciprocally connecting to one another more sporadically. Moreover, long-range IG*-Vglut1* projections link limbic to olfactory circuits. Lastly, we generated *Scgn*^-/-^ mice, in which IG*-Vglut1* neurons exhibited reduced intrinsic excitability. At the organismal level, the lack of *Scgn* triggered neuronal sensitization to methamphetamine (*Meth*) in the form of epileptic seizures, a major adverse condition experienced by adolescent Meth users (22, 23).

## Results

### Brain-wide activity mapping of prenatal drug exposure

Even though detailed knowledge using immediate early genes as cellular marks reflecting neuronal activation in response to psychoactive drugs exists (24–27), brain-wide and comparative activity mapping of psychostimulant drug effects in the developing and adult brains has not been performed to date. Here, we induced *Fos-*CreER^T2^::*ZsGreen1*^stop-floxed/stop-floxed^ dams (15) with tamoxifen on E14.5. This was 24h prior to administering amphetamine (10 mg/kg) (28, 29), nicotine (2 mg/kg) (11) or caffeine (6 mg/kg) (30), the most widely-used psychostimulants in general (31) and during pregnancy (32), on E15.5 with 48h survival (Fig. 1A; Fig. S1A) to allow the sufficient loading of tamoxifen-primed, drug-activated cells by *ZsGreen1* (Fig. S1B,C). By using c-Fos histochemistry, we confirmed the efficiency and specificity of Fos-driven *Cre*-mediated recombination in the fetal cerebrum (Fig. 1B). Thereafter, reconstructive light-sheet microscopy for brain-wide cell activation at E17.5 (Videos S1-4) showed that all drugs increased *ZsGreen1* expression relative to vehicle-treated controls (Fig. 1C). *ZsGreen1* was seen in fetal and adult neurons and perivascular (endothelial) cells (Fig. S1D) but neither astroglia nor micro-/oligodendroglia (Fig. 1D,E), even though a subclass of astroglia (‘immediate-early astrocytes’) can express *c-Fos* in response to glutamate and inflammatory stimuli (26, 33). The preferential labeling of neurons in the fetal brain is also compatible with the late-onset of astrogliogenesis, peaking neonatally, in rodents (34).

**Figure 1:**
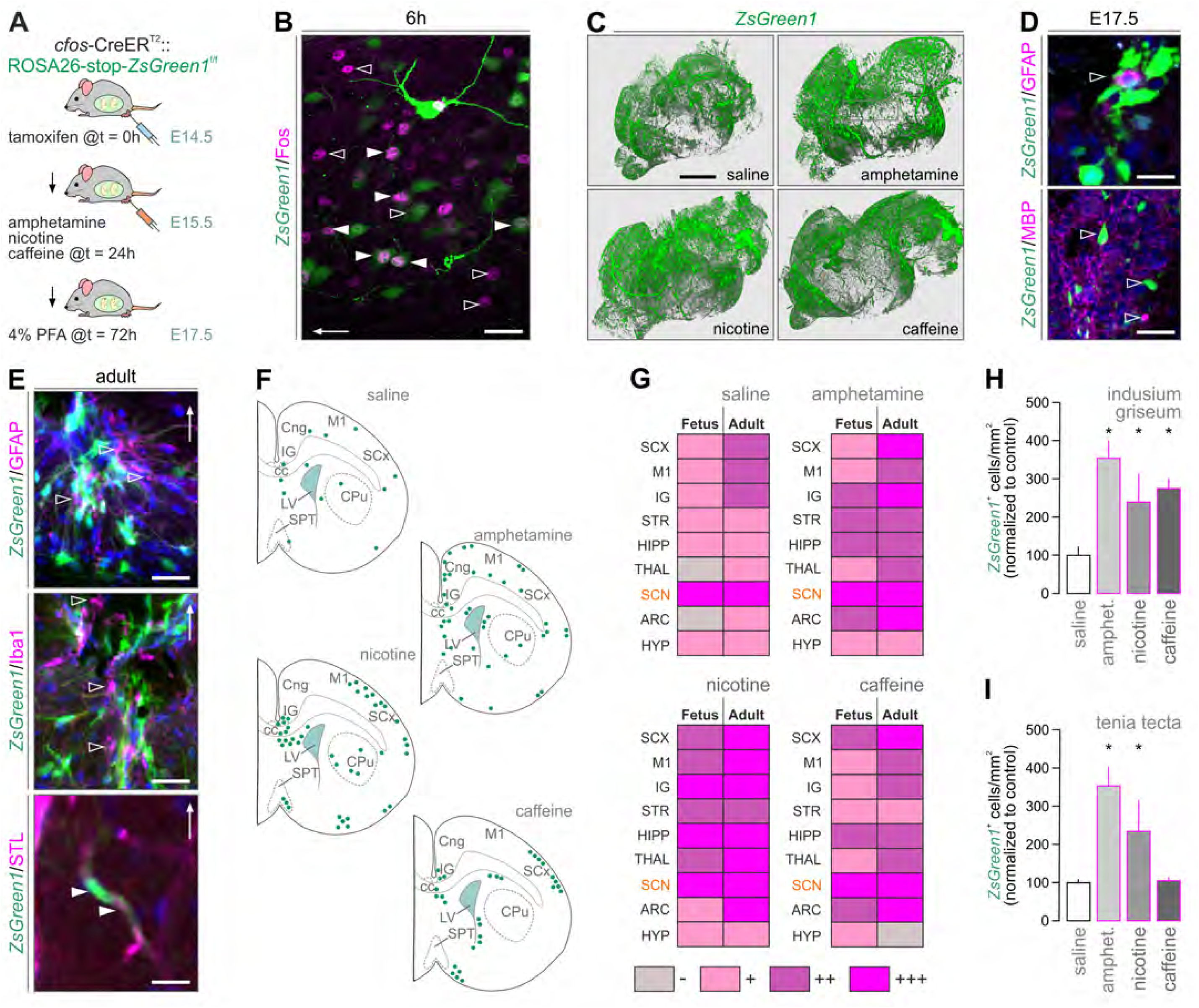
Brain-wide and trans-generational cell activation maps upon psychostimulant exposure. (**A**) Experimental design in *Cfos*-CreER^T2^::*ZsGreen1*^stop-floxed^ dams for the correlated analysis of fetal and adult brains. (**B**) *ZsGreen1*^+^ neurons label for c-Fos (*solid arrowheads*) 6h after amphetamine injection. The presence of only *ZsGreen1*^+^ neurons *(open arrowheads*) is taken as an indication of biologically-relevant and transient c-Fos expression. A cortical area is shown; arrow indicates the pial surface. (**C**) 3D rendering of optically-cleared fetal *ZsGreen1*^+^ brains after psychostimulant exposure. Neither fetal (**D**) nor adult (**E**) brains showed *ZsGreen1* signal in astroglia (GFAP^+^), microglia (Iba1^+^) or oligodendrocyte-like (MBP^+^) cells (*open arrowheads*). Endothelial cells (*arrowheads*) were visualized using *Solanum tuberosum* lectin (STL). In E, the IG is shown with arrows pointing dorsally. (**F**) Cellular *ZsGreen1* distribution in coronal sections spanning the IG. For abbreviations and consecutive levels see *Figure S3*. **G**) Comparison of cell activation patterns between fetuses and their respective dams (‘adult’). The suprachiasmatic nucleus (SCN) with its periodic *Fos* expression served as positive control. (**H,I**) Quantitative increases in *ZsGreen1*^+^ cell numbers in select brain areas upon psychostimulant treatment. Data were acquired using Arivis after light-sheet microscopy (C,H,I). (*Scale bars* = 1.5 mm (C), 30 μm (D *lower*), 20 μm (B,E), 10 μm (D *upper*).

*ZsGreen1*^+^ activity mapping in serial coronal sections of the fetal forebrain from saline-injected dams at E17.5 (Fig. 1F and S2,3) revealed basal *Fos*-CreER^T2^ activity in motor and sensory cortices, hippocampus proper, suprachiasmatic nucleus (SCN) of the hypothalamus. Considering that pacemaker systems of the SCN, including both neurons and astroglia (33, 35), can diurnally up-regulate *Fos* mRNA expression, we interpreted *ZsGreen1* labeling in this hypothalamic area as an innate biological control, particularly because of the uniform (treatment-independent) size of the *ZsGreen1*^+^ cell cohort (suggesting a ceiling effect due to rhythmic transcriptional activity; Fig. 1G and S3,4). All drugs induced *ZsGreen1* expression in the ventral tegmental area (Fig. S2), which we have taken as another positive control, showing the drug-induced activation of the reward circuitry. Specifically, amphetamine increased the number of *ZsGreen1^+^* cells in cingulate, motor and somatosensory cortices, IG (Fig. 1G,H), tenia tecta (TT; Fig. 1I), striatum, hippocampus, choroid plexus and ARC (Fig. S3). Nicotine induced *ZsGreen1* expression in cingulate, motor and pyriform cortices, medial septum (which harbors cholinergic neurons expressing nicotinic acetylcholine receptors) (36), hippocampus, MPN, ARC and supraoptic nucleus (Fig. S3). Besides substantial overlap with the effects of the other dugs, caffeine triggered notable *ZsGreen1* expression in the lateral diagonal band of Broca (27), MPN and ARC (25) (Fig. 1F,G) but not TT (Fig. 1I). These data demonstrate that prenatal psychostimulant exposure drives widespread and drug-specific neuronal and endothelial activation with the cingulate cortex/IG identified as a convergence point of drug action.

Next, we have repeated *ZsGreen1* mapping in the dams to characterize regional differences, if any, between fetal and adult brains. We have found significant psychostimulant-induced cell activation in cortical, thalamic and hypothalamic areas of which the cingulate cortex/IG and ARC invariably contained *ZsGreen*^+^ cells (Fig. S4). Thereby, we were able to reliably capture, using heat map-based checkerboard analysis (Fig. 1G), regional and age-associated differences in psychostimulant sensitivity in fetal *vs*. adult brains.

### Neuronal targets of psychostimulant action in the fetal and adult IG

Neuroanatomical studies separate the IG from the overlaying anterior cingulate cortex (ACA) in adult brain by it being negative for calbindin-D28k and, conversely, positive for *Scgn* or neurotrophin-3 (20, 37). Earlier studies suggest that *Scgn*, an EF-hand Ca^2+^-sensor protein evolutionarily linked to calbindin-D28k and calretinin, marks a prominent group of IG neurons whose biophysical signature resembles that of dentate granule cells (20). Therefore, *Scgn* is appealing as a marker of putative excitatory IG neurons. In addition, the IG receives dense monoaminergic and GABAergic innervation (38, 39). Its particularly prominent monoaminergic afferentation (Fig. S5A,B) that sets it apart from other cortical areas by mid-gestation might be a reason for its sensitivity to psychostimulant-induced perturbations.

Immunofluorescence histochemistry in combination with *Fos-*CreER^T2^-based genetic probing suggested that *ZsGreen1*^+^ neurons in the fetal IG could arise from the cortical ventricular zone: we have seen many fusiform *ZsGreen1*^+^ neurons with ramifying local processes in the corpus callosum and entering the IG and lining its callosal (ventral) surface (Fig. 2A). At E17.5, many *ZsGreen1*^+^ neurons were immunoreactive for the POU Class 3 Homeobox 3 (*Pou3f3*/*Brn1*) transcription factor (Fig. 2A), which broadly marks glutamatergic neurons, particularly pyramidal cells destined to cortical layer 2-4(40). Notably, none of the neurons populating the prospective IG expressed *Scgn* at E17.5, irrespective of them being eventually *ZsGreen1*^+^. This unexpected feature distinguishes the IG from other forebrain areas in which *Scgn* expression usually precedes that of other neuron-specific Ca^2+^-binding proteins (20). The lack of *Scgn* together with preferential D1 dopamine receptor expression (Fig. S5A) in callosum-associated glutamatergic neurons could explain, at least in part, their preferential psychostimulant-sensitivity in the fetal IG. This hypothesis is also supported by the finding that in the adult brain *ZsGreen1*^+^ neurons in the IG are small-to-medium in diameter, multipolar, *Scgn*^-^ and positioned relatively distant from the pial surface (Fig. 2B). These cellular features suggest non-excitatory identity. The fact that *Scgn*^+^ glutamatergic neurons were invariably *ZsGreen1*^-^ in the IG of the dams corroborates earlier observations on neuroprotection mediated by Ca^2+^-binding and sensor proteins (41–46) *Scgn/ZsGreen1* co-existence in the ARC served as positive (biological) control (Fig. 2B and S1D,E), which is compatible with their expression of nicotinic acetylcholine receptors and cocaine and amphetamine-regulated transcripts (47). Here, we have specifically mapped the developmental trajectory and psychostimulant sensitivity of *Scgn*^+^ excitatory IG neurons.

**Figure 2:**
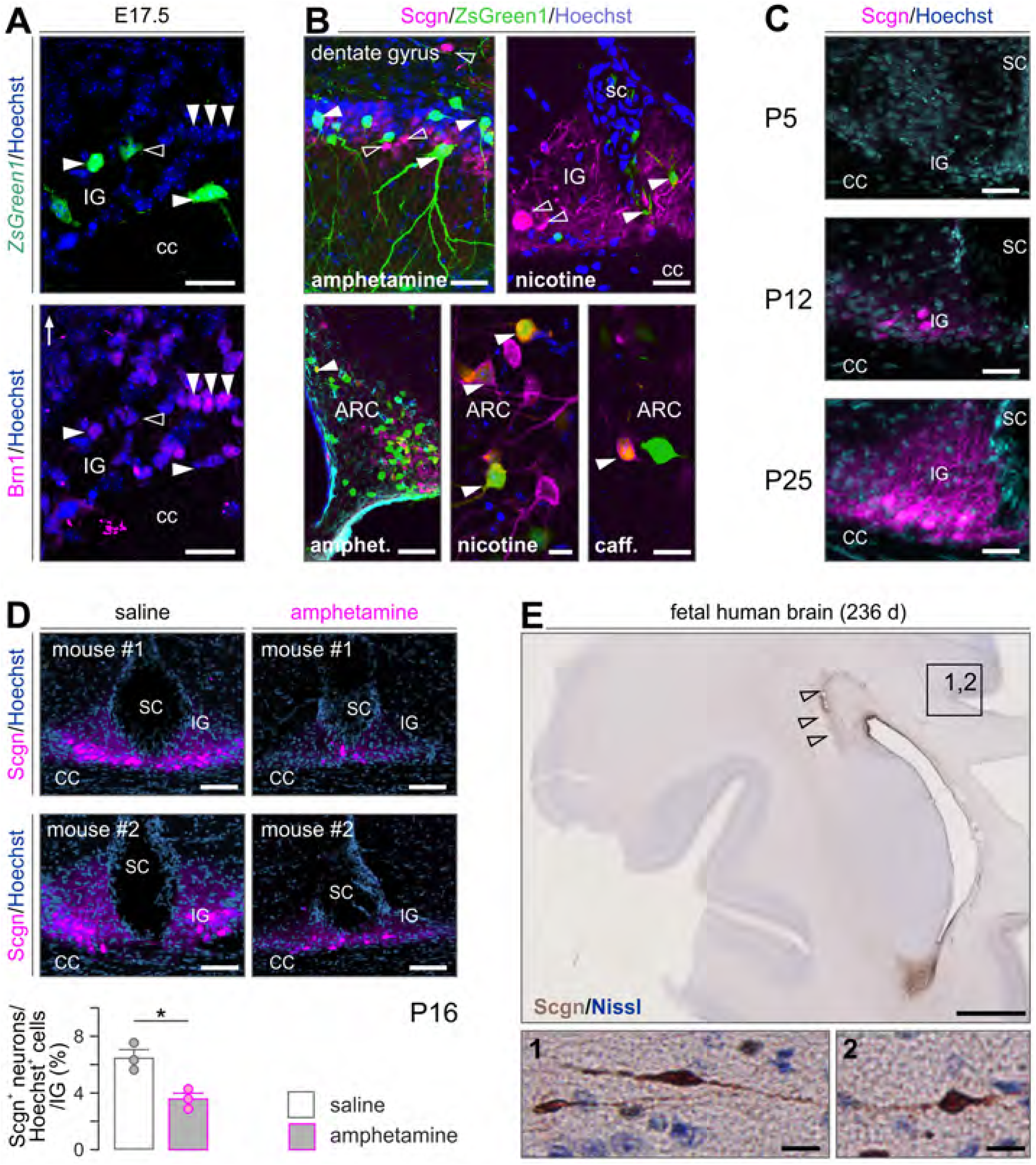
Psychostimulant sensitivity and ontogeny of *Scgn*^+^ neurons in the indusium griseum. (**A**) At E17.5, many psychostimulant-responder neurons in the indusium griseum (IG) express *Pou3f3/Brn1* (*arrowheads*; open arrowhead shows the lack of co-localization), a transcription factor for cortical pyramidal cells(40), and coincidently lack *Scgn*.(**B**) In contrast, in the adult dentate gyrus, indusium griseum (IG) and anterior hippocampal continuation (*top row*), psychostimulant-induced *ZsGreen1* expression (*solid arrowheads*) did not label *Scgn*^+^ neurons (*open arrowheads*). Yet *Scgn*^+^ neurons of the arcuate nucleus of the hypothalamus (ARC) showed coincident *ZsGreen1* expression (*solid arrowheads*). (**C**) *Scgn* localization in the mouse IG during postnatal development with postnatal day (P)12 taken as the earliest time point at which *Scgn* was successfully detected. (**D**) Prenatal amphetamine exposure significantly reduces *Scgn* expression in the IG on P16; *p* < 0.05, *n* = 3/genotype. (**E**) *Scgn*^+^ neurons line the IG in the human fetal brain. *Top*: Coronal section of a 236-days old human fetal brain at the level of the anterior horn of the lateral ventricle. Arrowheads indicate *Scgn* expression in the rostral migratory stream. *1,2*: *Scgn*^+^ neuroblasts in the lateral part of the IG, on the dorsal surface of the genu/truncus corporis callosi. *Abbreviations*: caff., caffeine; cc, corpus callosum; sc, sulcus centralis. *Scale bars* = 1 cm (A), 120 μm (B *bottom left*,D), 50 μm (B *top left*), 30 μm (A,B *top right*,C), 12 μm (B *bottom middle, right*, E/1,2).

First, we find *Scgn* be gradually enriched in the IG from postnatal day (P) 12 and reaching adult levels by P25 (Fig. 2C). Likewise, neuropeptide Y expression was transient in IG and detectable only on P5 (Fig. S5B-D). These observations suggest that excitatory neuron development is a protracted process in the IG, likely limiting the excitability of this neuronal contingent during fetal development. Next, we argued that intrauterine psychostimulant exposure might perturb *Scgn* expression in IG neurons if Scgn itself is a critical and activity-dependent differentiation mark. To test this hypothesis, we have treated pregnant mice with amphetamine (10 mg/kg) as above and sampled their male offspring on P16, a time point when *Scgn* level is substantial yet sub-maximal (Fig. 2D). As such, prenatal amphetamine application resulted in a significantly delayed Scgn expression in the IG (Fig. 2D). These data directly link *Scgn* expression to psychostimulant sensitivity in excitatory neuronal progenies of the developing IG.

Considering that neither cellular nor detailed network features of *Scgn*^+^ IG neurons are known (beyond some rudimentary biophysical characteristics (20)), we have molecularly characterized *Scgn*^+^ IG neurons using the *Patch-seq* workflow (48). Subsequently, we addressed if genetic ablation of *Scgn* increases neuronal sensitivity to methamphetamine.

### Scgn marks the mouse and human IG

The supracallosal domain of the IG resides in two parallel narrow stripes of grey matter situated rostro-caudally on the dorsal surface of the corpus callosum (CC) (Fig 3A,B). Once stretching over the genu of the CC, a small descending dorsoventral line of IG neurons forms the anterior hippocampal continuation (AHC) (Fig 3B). Earlier, we have shown that *Scgn* marks the IG and AHC, which share most microanatomical characteristics, in both lower primates and rodents (20). These data are recapitulated here (Fig. 3A-F) with a novel antibody (*see* Methods; Fig. S5F), and extended by CUBIC-based tissue clearing and intact tissue imaging to show that *Scgn* also labels the axonal bundle of the IG and AHC turning under the genu of the CC (Fig 3G). Moreover, our localization of *Scgn*^+^ neurons in the pyramidal layer of the CA1 and CA2 subfields of the mouse hippocampus (Fig. 3C), as well as the dentate gyrus (DG) and FC, visualizing the extrahippocampal formation (Fig. 3C), reconciles data from *post-mortem* human hippocampal CA1, where *Scgn* was implicated in conferring resistance to neurodegeneration (49).

**Figure 3:**
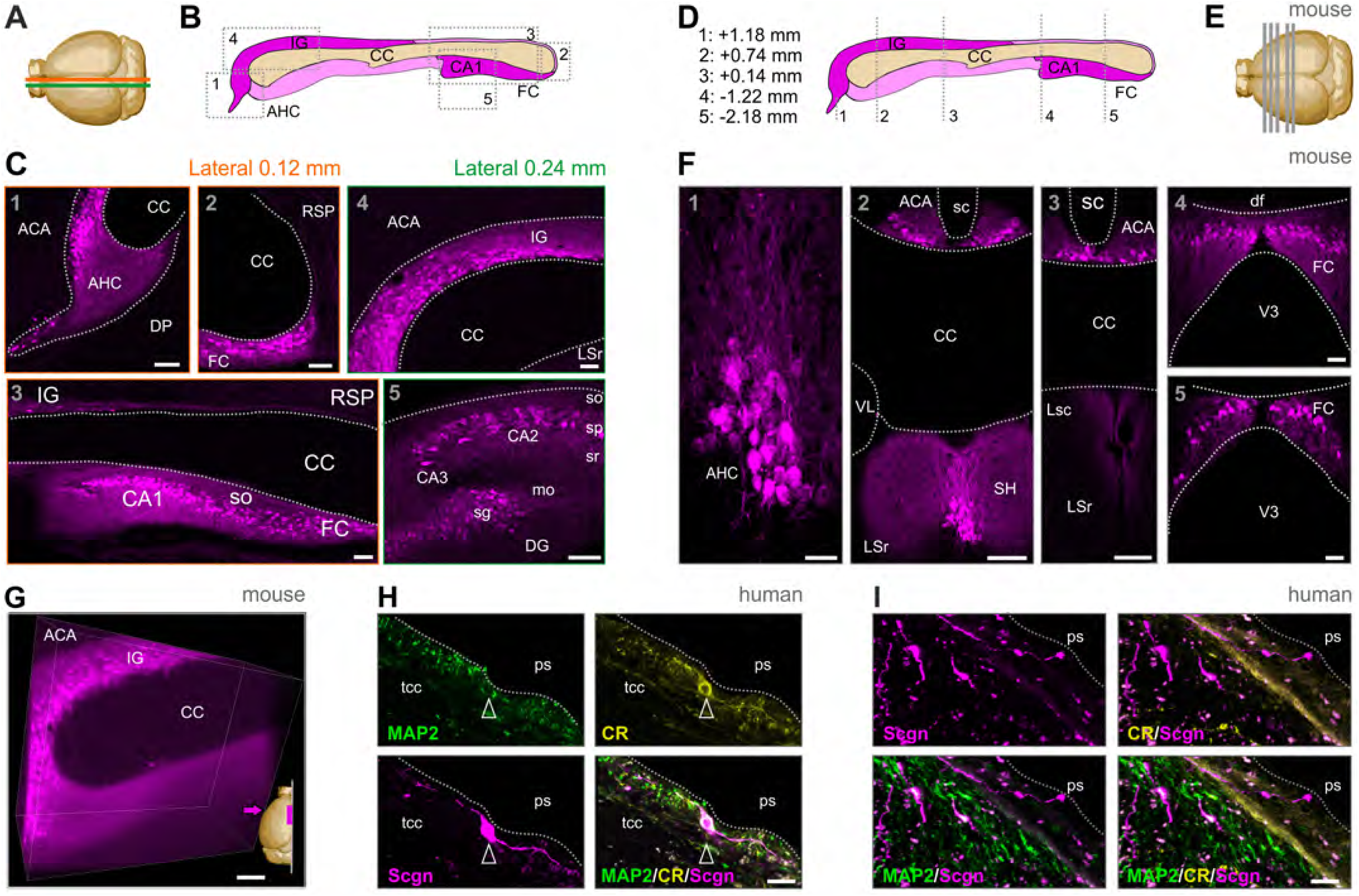
Anatomy of the mouse and human indusium griseum. (**A**) Brain scheme with orange and green lines indicating the cutting planes at 0.12 mm and 0.24 mm from the midline. (**B**) Sagittal scheme of the mouse extra-hippocampal formation, including IG. Grey dashed rectangles indicate the location of panels in C. (**C**) *Scgn* immunoreactivity (magenta) in the anatomical regions of the extra-hippocampal formation defined by the Allen brain atlas. (**D**) Coronal scheme of the mouse extra-hippocampal formation. Grey dashed lines (equally spaced for illustration) and coordinates relative to bregma indicate the coronal cutting planes shown E. (**E**) Brain scheme with coronal planes of tissue sampling. (**F**) *Scgn* immunoreactivity at the anterior-posterior coordinates indicated. (**G**) 3D image of optically-cleared IG with *Scgn* immunostaining. A 500-μm sagittal block is shown. Magenta arrow indicates the point of view. (**H**) Human IG with a *Scgn*^+^ IG neuron close to the pial surface (ps) expressing calretinin (CR) but not MAP2. (**I**) *Scgn*^+^/CR^-^ bipolar neurons (*arrowheads*) in human IG. Both *Scgn*^+^/MAP2^+^ and Scgn^+^/MAP2^-^ neurons could be found. *Abbreviations*: ACA, anterior cingulate area; AHC, anterior hippocampal continuation; CA1-3, Cornu Ammonis 1-3 subfields; cc, corpus callosum; df, dorsal fornix; DG, dentate gyrus; DP, dorsal peduncular cortex; FC, fasciola cinerea; LSc/r, lateral septal nucleus caudal/rostroventral; mo, straum moleculare; RSP, retrospinal area (ventral); sc, sulcus centralis; SH, septohippocampal nucleus; sg, so, straum oriens; sp, stratum pyramidale; sr, stratum radiatum; tcc, truncus corporis callosi; VL, lateral ventricle; V3, third ventricle. *Scale bars* = 100 μm (F/2-4), 50 μm (C/1-5,E,F/4,G), 20 μm (F/1,5), 10 μm (H,E).

While the rodent and primate IG are known to contain neurons (20), the cellular organization of the human IG is considered different with glial cells recognized as its building blocks (21). This notion prevails despite the fact that, at the gross anatomy level, the IG in humans is reminiscent of its mouse equivalent by being adherent to the lamina terminalis and extending between the genu and splenium of the CC. When using *Scgn* as a molecular marker (Fig. 3H,I), we find bipolar neuron-like cells with fusiform perikarya and long processes in both the fetal (3^rd^ trimester; Fig. 2E) and adult human IG (Fig. 3H,I). The neuronal identity of these cells is supported by their expression of either calretinin (Fig. 3H) or microtubule-associated protein 2 (MAP2; Fig. 3I), even though these markers seem abundant in non-overlapping cell cohorts. Thus, we suggest that *Scgn*^+^ neurons with morphological features reminiscent of relay or dormant migratory neurons earlier found in the olfactory tubercule of humans (45) also exist in the human IG.

### Long-range connectivity of *Scgn*^+^ FC/IG neurons

The extra-hippocampal pathway with its continuum of *Scgn*^+^ neurons is a complex integration loop caudally exiting the hippocampus through the subiculum towards the FC. To map out if long-range connections exist between the FC and IG, biotinylated dextran amine (BDA) was injected into the FC and used to anterogradely trace FC efferents (Fig S6A). BDA^+^ axons travelled around the splenium of the CC and split into parallel tracts at its ventral and dorsal surfaces (Fig S6B). BDA^+^ axons terminated on the dendrites of *Scgn*^+^ IG neurons (Fig S6C,D) and eventually ran parallel with the dendrites of *Scgn*^+^ neurons (Fig S6E). Next, we have tested if *Scgn*^+^ neurons receive synaptic inputs from the olfactory bulb, an experiment prompted by the notion that *Scgn*^+^ neurons also line the rostral migratory stream (as “shell cells”) (45) and populate the olfactory bulb. Here, cholera toxin B was used as retrograde tracer and injected into the main olfactory bulb. Indeed, populations of *Scgn*^+^ neurons along the IG were seen co-labeled for cholera toxin B and *Scgn* (Fig. S6F). These anatomical features suggest that *Scgn*^+^ neurons can relay information along the FC-IG-olfactory bulb circuit (Fig. 4A).

**Figure 4:**
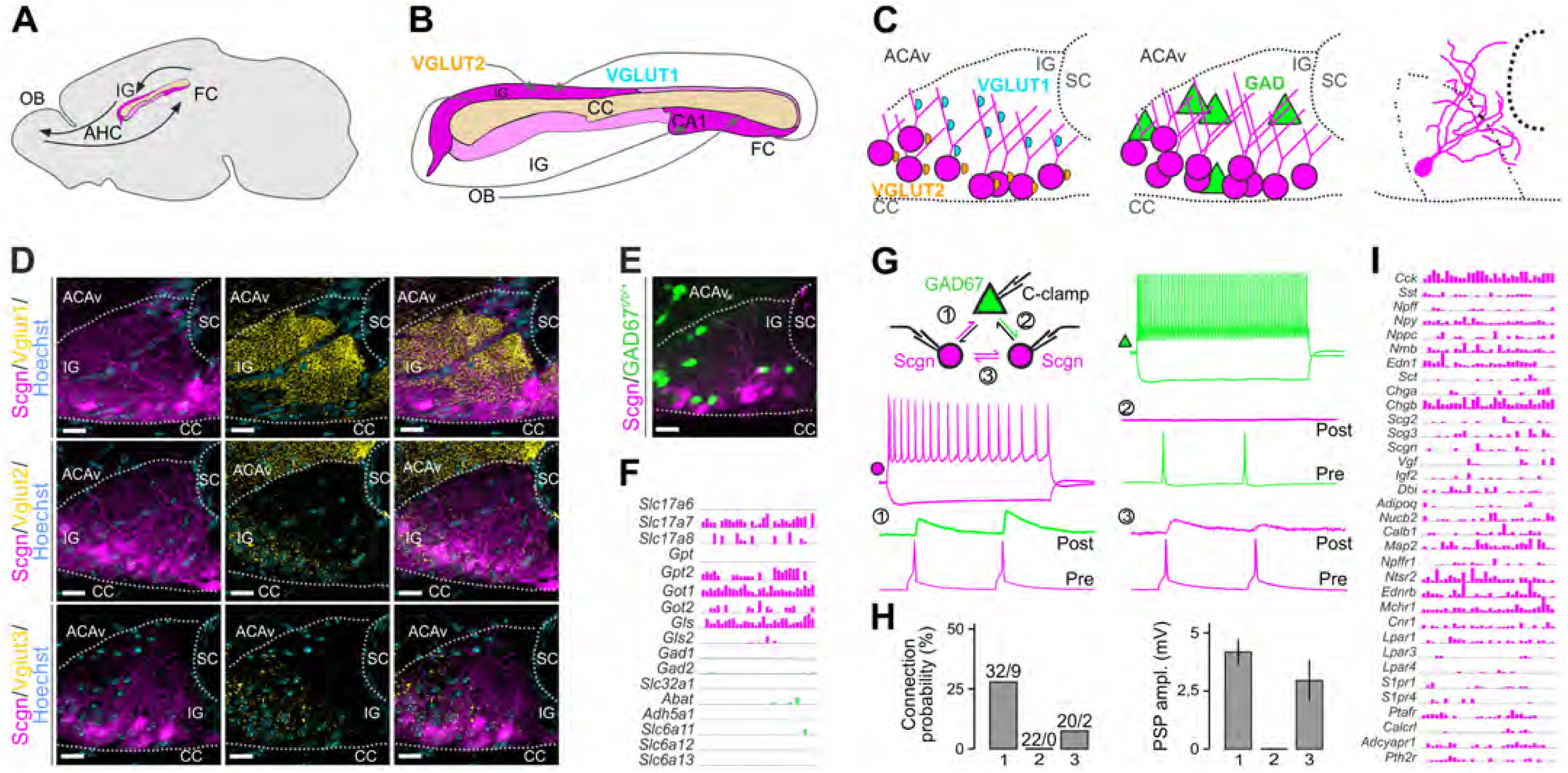
Local networks in the indusium griseum. (**A**) Sagittal scheme of the mouse extrahippocampal formation with the CC and IG and putative connectivity between the olfactory bulb (OB) and fasciola cinerea (FC) indicated. (**B**) Predicted glutamatergic connections, including the integration of excitatory afferents containing VGLUT1 and VGLUT2. (**C**) Coronal scheme of the IG with the cellular positions of VGLUT1^+^ and VGLUT2^+^ synaptic puncta (l*eft*), the location of excitatory *Scgn*^+^ *vs.* GAD67^gfp/+^ GABAergic neurons (*middle*), and reconstruction of a biocytin-filled *Scgn*^+^ neuron (*right*). (**D**) Distribution of VGLUT1, VGLUT2 and VGLUT3 in the IG. Note the predominance of VGLUT1^+^ innervation. Sections were at the level of +0.6 mm from bregma. (**E**) Spatial distribution of excitatory *Scgn*^+^ and GAD67^gfp/+^ GABAergic neurons in IG. (**F**) Expression of neurotransmitter modality-related genes identified with *Patch-Seq*. Log2 expression of genes coding for glutamatergic (magenta) and GABAergic (green) neurotransmitter synthesis, vesicular loading and degradation. Numbers indicate possible connected neurons. (**G**) *Top left*: Setting of *ex vivo* triple whole-cell electrophysiology recordings. *Middle*: Representative firing patterns of *Scgn*^+^ (magenta) and GAD67^gfp/+^ (green) neurons (top), with rheobasic spikes in grey. 1,2 and 3 shows combination of possible connections with representative examples of paired recordings. (**H**) Bar graphs depict the connection probability (*bottom left*) ad amplitudes of recorded PSPs (*bottom right*). (**I**) Neuropeptide and neuropeptide receptor gene expression in IG neurons, revealed by *Patch-seq*. *Abbreviations*: ACAv, anterior cingulate area (ventral); AHC, anterior hippocampal extension; CA1, Cornu Ammonis sub-filed 1; cc, corpus callosum; sc, sulcus centralis.

### Synaptic inputs to and local circuits of *Scgn*^+^ IG neurons

Neuronal computation in the IG summates the activity of subcortical cholinergic and midbrain monoaminergic territories, which is supported by the accumulation of afferent inputs harboring choline-acetyltransferase (ChAT), dopamine β-hydroxilase (DBH), tyrosine hydroxylase (TH), D2 dopamine receptors (Fig. S5B) or 5-HT receptors (5-HTR7) (38, 39). This anatomical complexity suggests that IG neurons might read out source network information for feedback to the hippocampus. The diversity of inputs is testable by e.g. histochemistry given the dichotomy of vesicular glutamate transporters (*Vgluts*) with *Vglut1* and *Vglut2* segregating to cortical and thalamic sources (Fig. 4B), respectively (50, 51). We find *Vglut1*^+^ or *Vglut2*^+^ synaptic terminals on *Scgn*^+^ IG neurons organized by a strict topography (Fig 4C,D) with *Vglut1*^+^ boutons targeting dendritic compartments, whereas *Vglut2*^+^ terminals arriving to neuronal somata (Fig. 4C,D). *Vglut3*^+^ terminals were found scattered along the somatodendritic axis (Fig. 4D). GABAergic interneurons populate the dorsolateral domain of the IG (Fig. 4C,E), and are recruited to local circuits. The lack of co-localization between *Scgn* and GFP in GAD67^gfp/+^ mice in intact, CUBIC-cleared tissues (52) reinforces that large-diamater *Scgn*^+^ IG neurons are non-GABAergic (Fig. 4E); instead are entrained by local GABA sources.

Besides their macrocircuit constellation, little is known about the local microcircuitry in which *Scgn*^+^ IG neurons are embedded into. Hence, we probed local connectivity using triple whole-cell patch-clamp recordings in GAD67^gfp/+^ mice (Fig. 4G and S7), where we visually distinguished fast-spiking (FS) GABA interneurons and GFP^-^ IG neurons, the latter being RS (Fig. 4G). We used triples consisting of 1 GAD67^gfp/+^ and 2 GFP^-^/RS IG neurons to test if excitatory IG neurons (*post-hoc* identified as *Scgn*^+^) are densely interconnected for local amplification and summation of neural information. We used the electrophysiological profile described by Mulder *et al*. (20) to identify *Scgn*^+^ neurons. This setting was also advantageous to assess reciprocal connectivity of *Scgn*^-^/FS and RS IG neurons (tested 72 putative connections; Fig 4G,H). We found ∼27% connection probability from RS-to-FS neurons (9 out of 32 tested; Fig 4H) with an average amplitude of 4.1 ± 0.7 mV. Reciprocal connections from FS-to-RS neurons were not found. Determination of *Scgn*^+^ IG neuron interconnectivity revealed the presence of excitatory connections in ∼10% of the cases (2 out of 20; Fig 4H), with an amplitude of 2.7 ± 1.1 mV. Thereby, we hypothesize that activation of the excitatory component of the IG is primarily achieved by cortical and subcortical synaptic inputs (38, 39) rather than through self-amplification loops locally.

### *Patch-seq* profiling of Scgn^+^ neurons

The only existing dataset on *Scgn*^+^ neurons posits similarities between DG and IG neurons on the basis of biophysical similarities, which is compatible with a broader view on the IG being an anterior extension of the DG (53). However, molecular analysis is required for this hypothesis to stand. Therefore, we have fed RNA samples from all neurons that had undergone electrophysiological recordings into our *Patch-seq* workflow (48) to achieve precise gene expression profiling (Fig. 4F,I and S8).

The analysis of genes associated with inhibitory or excitatory neurotransmission in RNA aspirates of IG neurons revealed the predominance of glutamate synthetizing and degrading enzymes (Fig. 4F). In contrast, the enzymatic machinery for GABA synthesis (e.g. *Gad1*, *Gad2*) was not expressed (*detection threshold* = >2 counts in >5/30 IG neurons) (Fig 4F). To use precise molecular mapping to predict (and support) how IG neurons are synaptically wired in the IG, we extracted single-cell RNA-seq data for neuropeptides and receptors (Fig. 4I). We find *Scgn* as a marker gene at low transcript levels, an observation compatible with its slow protein turnover (54). Therefore, homogeneity of the electrophysiological signatures <u>and</u> molecular parameters of non-GAD67^gfp/+^ IG neurons was used as cellular classifier (Fig. 4I). Excitatory IG neurons expressed endothelin receptor type B (*Ednrb*), neurotensin receptor 2 (*Ntsr2*), melanin-concentrating hormone receptor 1 (*Mchr1*), cannabinoid receptor 1 (*Cnr1*) and parathyroid hormone receptor 2 (*Pthr2*) indicating the physiological role of these neurons in distinct hormonal responses, embedding IG neurons into hormonal feedback networks. In addition, mRNA transcripts for many classical and novel neuropeptides were also seen, including cholecystokinin (*Cck*), somatostatin (*Sst*); neuropeptide FF (*Npff*), neuropeptide Y (*Npy*), natriuretic peptide C (*Npcc*), neuromedin B (*Nmb*), chromogranin B/secretogranin B (*Chgb*) and calbindin D28k (*Calb1* at detection threshold = >2 counts in >5/30 IG neurons; Fig 4I).

Even though mRNA expression is taken broadly indicative of neuronal identity, mRNA-to-protein translation does not necessarily occur, which can bias the biological interpretation of single-cell RNA-seq data. Here, we show that despite the presence of *Npy* mRNA in all the *Patch-Seq*-recorded neurons, NPY immunoreactivity was not detectable in adult IG. Instead, we find NPY-expressing neurons at P5 (Fig. S5C), indicating a putative and transient developmental role for NPY, equivalent to other neuropeptides in the neonatal period (55). Similarly, the presence of CCK mRNAs in IG was reported in our earlier work on a CCK^BAC/DsRed^::GAD67^gfp/+^ composite mouse line (18, 48) even though conventional histochemistry fails to visualize CCK protein. Furthermore, lower levels of *Drd1* mRNA was also detected (*data not shown*) and confirmed by genetic labeling in the corresponding D1R-GFP mouse line (Fig. S5A).

Finally, we aimed to identify cell type-specific markers of excitatory IG neurons by using a contrast analysis between our dataset and cortical single-cell RNA-seq data (56). We filtered the set of cortical genes with zero or low expression (low-pass filtered by the interquartile range of expression values, i.e. lowest 25% of the expressed genes identified by Tasic *et al.* (56)) and compared these to highly-expressed genes in IG from our *Patch-seq* dataset (high-pass filtered to extract the highest 25% of expressed genes). This intersectional analysis returned interleukin-16 (*Il-16*), aggrecan (*Acan*), FERM domain-containing protein 2 (*Frmpd1)*, hepatocyte growth factor (*Hgf*) and Ca^2+^-binding protein-7 (*Cabp7*) as preferentially expressed in excitatory IG neurons (*vs*. cortical ones; Fig. S8B). Interestingly we found neither *Prox1* nor *Porx2* expression, which identify both fetal and adult dentate granule cells (57), thus suggesting the molecular demarcation of IG neurons at least at specific segments of their transcriptional control. These data establish the precise molecular make-up of excitatory IG neurons, set them apart from their cortical counterparts, and allow testing of both neuronal connectivity and hormone signaling.

### Adult *Scgn*^-/-^ mice are sensitive to methamphetamine

Methamphetamine (*Meth*), a derivate of amphetamine, is known to obstruct neuronal excitability (58, 59). Therefore, we first confirmed that acute superfusion of *Meth* (50 µM) directly affects RS IG neuron excitability during *ex vivo* whole-cell current clamp recordings. Indeed, *Meth* suppressed action potential (AP) formation in IG neurons, leading to a maximum of a single AP on a stimulus step (Fig. 5A), which showed a smaller amplitude and significantly increased half-width (Fig. 5B). *Meth* effect was instantaneous, robust and reversible (Fig. S9A) and also occurred in *Scgn*^-/-^ slices (Fig. S9B).

**Figure 5:**
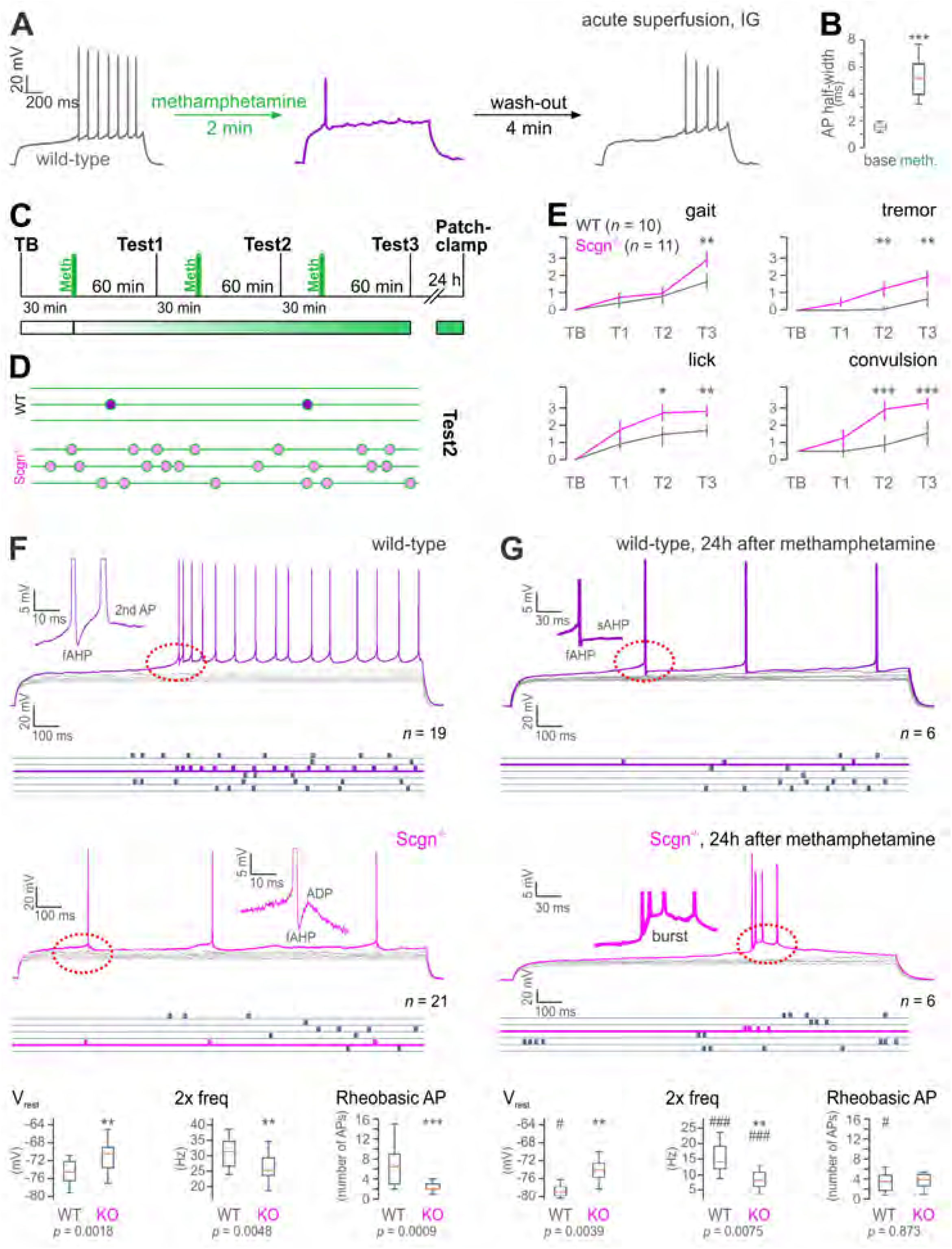
Scgn ablation sensitizes to methamphetamine. (**A**) Methamphetamine (*Meth*) superfusion acutely inhibits AP discharges in IG neurons in wild-type (WT) mice (*N* = 2/*n* = 4). (**B**) Significant AP half-width increase during bath application of *Meth*. (**C**) Experimental design of Meth administration, behavioral testing and *ex vivo* electrophysiology. (**D**) Convulsions in WT *vs*. *Scgn*^-/-^ mice during Test 2 (green lines represent 60 sec). (**E**) Summary of behavioral scores (gait, tremor, lick, convulsion) in TB and T1-T3 of Meth injected WT (*N* = 10) and *Scgn*^-/-^ mice (*N* = 11). (**F**) Excitability and intrinsic discharge properties of IG neurons from WT (*N* = 7/*n* = 19) and Scgn^-/-^ mice (*N* = 7/*n* = 21). Rheobasic responses in purple (WT) and pink (Scgn^-/-^) with raster plots showing firing rates in blue. Note the reduced firing rate, and the lack of a pre-potential (red dashed circle) in *Scgn*^-/-^ IG neurons (insets show ADP kinetics). Comparison of biophysical parameters from WT *vs*. *Scgn*^-/-^ neurons (Bottom). (**G**) Excitability and intrinsic properties of IG neurons from WT (*N* = 4/*n* = 6) and *Scgn*^-/-^ mice (*N* = 3/*n* = 6) 24 h after *Meth* exposure Note the reduced firing rate and decreased ADP in WT, as well as bursting in *Scgn*^-/-^ IG neurons (red dashed circles). (*Bottom*) Comparison of intrinsic parameters from WT *vs*. *Scgn*^-/-^ neurons 24 h after *Meth* exposure and their statistical comparison to naive state measures (**F;** labeled with #). *^/#^*p* < 0.05, **^/#^*p* < 0.01, ***^/###^*p* < 0.001; two-samples *t*-test. *Abbreviations*: ADP, after-depolarization; AP, action potential; APthr, AP threshold; TB, test at baseline; T1-T3, tests 1-3; Vrest, resting membrane potential; 2x freq, firing frequency at 2x rheobase stimulation.

High-dose or repeated *Meth* administration triggers extensive neuronal apoptosis and astrogliosis in the IG *in vivo* (28, 29, 60). As a behavioral effect downstream, *Meth* can either trigger epileptic seizure-like activity (23, 61, 62) or enhance those if pre-exist. Given the sensitivity of *Scgn*^+^ limbic (including IG) neurons to psychostimulants, we asked if genetic deletion of this Ca^2+^-sensor would alter the sensitivity of the IG circuit to *Meth*; demonstrated by increased seizure activity. This hypothesis rests on the knowledge that the loss of other Ca^2+^-sensors and buffers perturbs neuronal excitability (44, 63, 64). Here, we injected both wild-type (*n* = 10) and *Scgn*^-/-^ mice (*n* = 11) with Meth (15 mg/kg, i.p., 3 times at 90 min intervals; Fig. 5C) and scoring animal behavior by the SHIRPA protocol (65, 66) 60 min after each Meth cycle (T1-3; Fig. 5C,E). Amongst the parameters scored, gait, tremor, licking behavior and clonic convulsions were significantly different between the genotypes with *Scgn*^-/-^ mice showed significant worsened signs towards Meth acutely (Fig. 5E). Clonic convulsions in *Scgn*^-/-^ mice at T2 was particularly notable (Fig. 5D). *Scgn*^-/-^ mice (54, 67) also produced more severe symptoms as fragmented locomotion and increased wall/floor licking, which often coincided with continuous reversing (Fig. 5E; Videos S5,6). These data show that *Scgn* ablation increases *Meth* vulnerability.

Next, we asked if *Scgn* is critical in determining pathophysiological changes in the biophysical characteristics of IG neurons by comparative analysis of wild-type *vs*. *Scgn*^-/-^ mice (54, 67) using whole-cell patch-clamp recordings. For wild-type IG neurons, we have identified the rheobasic AP threshold (APthr) by applying 1-2 pA incrementing current steps. All cells hosed a pre-potential when reaching the APthr and continued spiking thereafter (Fig. 5F, *top*). Herein, after-depolarization following a fast after-hyperpolarization phase was sufficient to evoke a 2^nd^ AP immediately after the 1^st^ rheobasic AP (Fig. 5F, *middle*). In contrast, *Scgn*^-/-^ neurons did not produce a pre-potential along with a reduced number of APs on their rheobasic responses (Fig. 5F and S10). Their ADP amplitudes were also significantly reduced. In sum, quantification of all AP parameters and firing patterns revealed significant differences between wild-type (*n* = 19) and *Scgn*^-/-^ (*n* = 21) RS IG neurons in, e.g., resting membrane potential (Vrest), number of APs on the rheobasic step and double-threshold firing frequency (2x Freq) (Fig 5F, *bottom*).

Finally, we sought to determine if *in vivo Meth* administration differentially affects the excitability of IG neurons in wild-type *vs*. *Scgn*^-/-^ mice. To this end, we performed patch-clamp recordings in *ex vivo* brain slices 24 h after behavioral testing (Fig. 5C). A history of *Meth* exposure significantly lowered the firing rate on rheobasic responses (to levels similar to those in *Scgn*^-/-^ neurons at baseline) and eliminated the ADP in wild-type IG neurons (Fig. 5G). In turn, *Scgn*^-/-^ IG neurons expressed a bursting rheobasic response (yet no change in frequency) 24h after *Meth* exposure, which appeared with considerable time lag upon rheobasic stimuli (Fig. 5G). *Post-hoc* morphological reconstruction confirmed that the same subtype of RS neurons at comparable locations in the ventromedial IG were probed (Fig. S10) and their lesser Meth responses upon Scgn knock-out (Fig. S11) to define the biophysical footprint of *Meth* administration. Thus, *Meth* is identified as a psychostimulant that retunes basic biophysical properties (e.g., Vrest) and capacity to fire AP trains in a *Scgn*-dependent manner (Fig. 5G). Cumulatively, these data provide a sufficiently broad set of indications to suggest that *Scgn* ablation compromises IG neurons in their level of excitability and sensitizes them to Meth-induced maladaptation, which might be relevant to scaling psychostimulant sensitivity along the DG→FC→IG→olfactory circuit.

## Discussion

Parcellation of cortical areas through the correlated molecular, cellular and circuit-level interrogation of their cellular, primarily neuronal, constituents revolutionized modern neurobiology by distinguishing a kaleidoscope of novel cellular subtypes (or modalities (68, 69)). However, studies which assign select neuronal contingents as function determinants to brain-state changes upon environmental stimuli (irrespective of these being adverse or benign) are at their infancy. Here, we combined psychostimulant-induced brain-wide genetic activity mapping (15) with *Patch-seq* interrogation of neurons (48) to capitalize on the advantages of the single-most powerful neuronal phenotyping techniques available to date. Our studies pinpoint the IG as the limbic area containing neurons ultimately activated by amphetamine (*Meth*), nicotine and caffeine. A plausible reason for such sensitivity is the unique developmental trajectory, molecular/neurochemical identity and circuit complexity of excitatory IG neurons; including the late postnatal expression of *Scgn*, a Ca^2+^-sensor protein. Moreover, the IG receives dense innervation from monoaminergic and cholinergic subcortical afferents (38, 39), which we view as a molecular underpinning for its selective sensitivity to psychostimulants. The neurochemical complexity of long-range afferentation together with the hierarchical distribution of glutamatergic afferents along the somatodendritic axis of excitatory IG neurons produces a strict organizational layout for the integration of synaptic inputs from cortical/hippocampal (*Vglut1*^+^; arriving to dendrites), subcortical/olfactory (*Vglut2*^+^; arriving to somata) and midbrain monoaminergic inputs (*Vglut3*^+^, TH^+^/DAT^+^ targeting perisomatic domains).

By textbook definition, the IG is referred to as a glia-laden area. This view is in part supported by developmental biology data from rodents and humans associating the IG with callosal development, which relies on glia-derived guidance cues (34, 70, 71): pioneer axons of the corpus callosum are spatially organized by the IG and the glial wedge dorsally, with the subcallosal sling positioned ventrally (72). This opinion is further escalated by the cellular anatomy of the human IG, which is perceived as a glial plate (21) probably due to the lack of neuronal markers, particularly those helpful to visualize fine neuronal morphology. The ‘glial dominance hypothesis’ persists despite detailed knowledge on the existence of immature neurons in the IG at birth, which reach mature neuronal morphology by P15 in rodents (including receptor expression) (70, 71).

Notably, the subcallosal sling was recently redefined as a neuron-enriched territory (72). Likewise, we make an effort to re-classify the IG as a neuronal domain given the existence of functionally-specialized and callosum-associated glutamatergic neurons to reside ventrally, while GABA cell populations accumulate dorsolaterally. By relying on *Scgn* and neuropeptides, we show that excitatory IG neurons are immature at P5, become neurochemically-specialized by P12-P16, by which time *Scgn* immunolocalization fully resolves the principal circuit design of both the IG and AHC. In fact, *Scgn* alone is sufficient to outline a DG→FC→IG→AHC→olfactory bulb loop, in which tract tracing distinguishes FC neurons to project to the IG that, in turn, innervate the olfactory bulb. Morphologically, *Scgn*^+^ IG neurons in rodent and human brains are similar, with uniquely long processes running parallel with the callosal surface. Our data on the existence of *Scgn*-only *vs*. *Scgn*^+^/calretinin^+^ neuronal subsets suggest essential heterogeneity or the activity-dependent regulation of *Scgn* (or calretinin) expression in human IG neurons. However, the use of *Patch-seq* itself carries sufficient precision to show that all *Scgn*^+^ neurons are glutamatergic, express e.g., *Cck*, *Npy*, *Adcyap1*, *Cnr1* and secretogranins (that is, neuropeptides, receptors and SNARE-components) and transcription factors (Fig. S8A) in unique constellations.

We then increased understanding on how the IG integrates synaptic information by studying local connectivity: local GABA interneurons receive significant innervation from *Scgn*^+^/*Vglut1*^+^ neurons. In contrast, *Scgn*^+^/*Vglut1*^+^ neurons themselves receive spatially-segregated glutamatergic innervation along their dendrites. Interestingly, excitatory IG neurons are only sparsely interconnected, highlighting GABA interneurons as critical circuit organizers. This model of circuit organization is compatible with the finding that non-Scgn neurons (likely interneurons) are primary drug responders in the adult IG, likely precipitating significant circuit breakdown by impairment of cellular “hubs” upon psychostimulant administration. The finding that IG neurons project to the olfactory bulb is significant when considering that they express receptors for endothelin, melanin-concentrating hormone, neurotensin and other hormones/neuropeptides, stressing that IG neurons differentially activated by a variety of brain-states could critically entrain olfactory circuits by delivering information antiparallel to olfactory output. Thus, a multimodal and reciprocal circuit emerges between olfactory and limbic areas.

Even though *Scgn* is a “pure” Ca^2+^-sensor given its low affinity to Ca^2+^ (73), its functional significance is only partially understood in, e.g., hypothalamic neurons (47). In the IG, it is conceivable to hypothesize that *Scgn* postnatally can protect against runaway excitation induced by psychostimulant exposure. This notion is supported by IG neurons being drug-sensitive when lacking *Scgn in utero*, and can have disease relevance because we consider *Scgn* expression to be activity dependent as supported by *i*) its distinct intracellular levels as a factor of localization (reflecting neuronal maturity) in the dentate gyrus and IG, and *ii*) limited mRNA detectability in *Patch-seq*-processed IG neurons (*n* = 8/22), which seems to reinforce the concept of basal expression before rapid use-dependent gene induction. Indeed, in *Scgn*^-/-^ mice we find reduced neuronal excitability and synaptic strength, as well as significant worsening of animal behaviors coincident with the progressive development of generalized (clonic) convulsive seizures upon exposure to *Meth in vivo.* These data link, for the first time, brain-wide activity maps to a specific molecular determinant of neuronal excitability that gates the brain’s vulnerability to illicit drugs. Thus, the IG is identified as a hub integrating (patho-)physiological information whose therapeutic resetting might alleviate adverse psychostimulant symptomatology.

## Author contributions

JF, TGMH and TH conceived the study. SR, AGM, FG, SK, GA, RAR, JH, KM and AA collected and analyzed data. YY and GGK contributed valuable tools and samples. TGMH and TH procured funding. JF and TH wrote the manuscript. All other authors commented on and approved submission of this manuscript.

The Authors declare no conflict of interest.

## Acknowledgement and funding

The Authors thank the Biomedical Sequencing Facility at the Center for Molecular Medicine of the Austrian Academy of Sciences, particularly T. Penz and C. Bock, for assistance with next-generation sequencing. Gilberto Fisone (Karolinska Institutet) is acknowledged for sharing Drd1-GFP and Drd2-GFP mice. This study was supported by the National Brain Research Program of Hungary (2017-1.2.1-NKP-2017-00002, AA), the Excellence Program for Higher Education of Hungary (FIKP-2018, AA), the Swedish Medical Research Council (TGMH, TH), Swedish Brain Foundation (‘Hjärnfonden’, TH), the European Research Council (ERC-2015-AdG-695136; TH) and intramural funds of the Medical University of Vienna (TH).

## Supplementary Information

### Materials and Methods

#### Animals and husbandry

We have generated a Scgn^tm1a(KOMP)Wtsi^ (*Scgn*^-/-^) mouse line which appeared anatomically normal, particularly without changes to its brain size or deformities to fine structures, including cell proliferation, migration and laminar distribution (1). We have also used GAD67^gfp/+^ knock-in mice to label GABAergic interneurons in the cerebral cortex and associated structures (2). *Drd1*-GFP (MGI:3840915) and *Drd2*-GFP (MGI:3843608) brains were from GENSAT. Wild-type mice refer to the C57Bl/6N strain. Animals were group-housed under 12:12 h/h light/dark cycle with *ad libitum* access to food and water. Mice of both sexes were used for neurophysiology experiments during the period of postnatal weeks 4-9, and for behavioral experiments during weeks 8-12. Experiments on live animals conformed to the 2010/63/EU European Communities Council Directive and were approved by the Austrian Ministry of Science and Research (66.009/0145-WF/II/3b/2014, and 66.009/0277-WF/V/3b/2017). Particular effort was directed towards minimizing the number of animals used and their suffering during experiments.

#### Tamoxifen injection and tissue processing

*Fos*-CreER^T2^::*ZsGreen1*^stop-floxed^ ‘TRAP’ dams (3) were injected with tamoxifen (10mg/kg) on embryonic day (E)14.5 to induce Cre-mediated recombination. Amphetamine (10 mg/kg), nicotine (2 mg/kg), caffeine (6 mg/kg) and saline (as vehicle) were injected on E15.5 with the animals sacrificed on E17.5 (48h activation; Fig. 1A). The brains of both the dams and their embryos were collected and immersion fixed in 4% PFA for 24h before being immersed into 30% sucrose for cryoprotection (48h). Adult brain tissues were cut on a cryostat as 50 µm-thick serial free-floating coronal sections. Embryonic brain tissues were cut as 14 μm thickness and mounted on fluorescence free glasses.

#### Prenatal exposure to amphetamine

On E15, pregnant C57BL/6N females (Charles River) were intraperitoneally injected with amphetamine (10 mg/kg body weight) or saline (as vehicle) at an injection volume of 10 ml/kg. All drugs were obtained from Sigma. Animals were kept separately under standard housing conditions with a 12h/12h light/dark cycle and food and water available ad libitum. On P16 and P20, offspring from each litter was randomly selected and sacrificed for analysis. Animals were fixed by transcardial perfusion with 50 ml of phosphate buffer followed by 100-150 ml of 4% paraformaldehyde (PFA) in phosphate buffer (PB, 0.1M, pH 7.4) and processed for histochemistry as described below.

#### Preparation of acute brain slices

All experiments were performed in 250 μm-thick para-coronal slices (Fig. S7) prepared on a VT1200S vibratome (Leica) in ice-cold artificial cerebrospinal fluid containing (in mM): 90 NaCl, 2.5 KCl, 1.25 Na2HPO4, 0.5 CaCl2, 8 MgSO4, 26 NaHCO3, 20 D-glucose, 10 4-(2-hydroxyethyl)-1-piperazineethanesulfonic acid (HEPES), 3 Na-pyruvate, 5 Na-ascorbate (pH 7.4). Brain slices were then incubated at 22-24 °C for 60 min in a recording solution containing (in mM): 124 NaCl, 2.5 KCl, 1.25 Na2HPO4, 2 CaCl2, 2 MgSO4, 26 NaHCO3, 10 D-glucose (pH 7.4). All constituents were from Sigma-Aldrich. Both solutions were aerated with carbogen (5% CO2/95% O2). Temperature in the recording chamber was set to 33 °C (TC-10, Npi). Brain slices were superfused with the recording solution at a rate of 4-6 ml/min. Neurons were visualized by differential interference contrast (DIC) microscopy on an Olympus BX51WI microscope.

#### Patch-clamp electrophysiology

Whole-cell recordings were carried out using borosilicate glass electrodes (Hilgenberg, Germany) of 3-4 MΩ pulled on a P-1000 instrument (Sutter). Electrodes were filled with an intracellular solution containing (in mM): 130 K-gluconate, 3 KCl, 4 ATP-Na2, 0.35 GTP-Na2, 8 phosphocreatine-Na2, 10 HEPES, 0.5 ethyleneglycol-bis(2-aminoethylether)- *N*,*N*,*N*′,*N*′-tetraacetate (EGTA), (pH 7.2 set with KOH), with Cl-reversal potential ∼-95 mV. Both single-cell patch-clamp recordings for cell harvesting and triple patch-clamp recordings were carried out on an EPC-10 triple amplifier (HEKA, Germany) controlled by PatchMaster 2.80. Current clamp recordings were corrected for −9.99 ± 0.38 mV liquid junction potential between the intracellular and recording solutions, as measured against a 3M KCl-electrode. Resting membrane potential (*V*rest, expressed as mVs) was measured in current clamp mode at 0 pA current upon breaking into whole-cell configuration. Input resistance (*Rm*, expressed as MΩ) was calculated using linear regression established between electrotonic voltage responses (±15 mV from *V*rest) and 500 ms current steps of increasing amplitude (10 pA increments). Membrane time constant (τ, ms) was averaged from 20 successive electrotonic voltage responses to hyperpolarizing (-40 pA) current steps and fitted with a single exponential. The sag depolarization, indicating the activation of a hyperpolarization-activated nonselective cationic current (*Ih*) was calculated as the ratio of the peak negative voltage and steady-state negative voltage at a voltage response to hyperpolarizing current injections resulting in a steady-state voltage of -100 mV. AP threshold (APthr, mV) was defined as the voltage point where the upstroke’s slope trajectory first reached 10 mV/ms. AP rise time (ms) was the time from the APthr to the AP’s peak. Maximum AP up- and AP down-strokes were determined as the maximum and minimum of the geometrical differential of the AP (mV/ms), respectively. Maximum up- and down-stroke times were the times from APthr to reach maximum AP up- and down-stroke, respectively. These parameters were measured for (*i*) the first AP elicited by a 1,000-ms rheobasic current step. Adaptation ratio was calculated as the ratio of the last 2 interspike interval relative to the first 2 interspike intervals. Firing frequency (Hz) was determined at double-threshold current injections producing spike trains. All parameters were measured by applying manual procedures custom-written in Matlab (MathWorks). We used 50 µM Meth (kind gift from Prof. Christian Pifl, CBR, Vienna) diluted in recording aCSF for acute Meth pharmacology probing of IG neurons. Histochemical visualization of increased neuronal activity after *in vivo* Meth administration experiments was carried out by c-fos immunolabeling using guinea pig anti-c-Fos (1,1:000, Synaptic Systems; Cat No. 226 005) in 1:500 concentration.

#### Cell harvesting for RNA sequencing

At the end of each patch-clamp routine, the micropipette was voltage-clamped to a holding potential (*V*hold) of -5 mV (4). For cell harvesting, the entire soma of each recorded neuron was aspirated into the micropipette slowly (∼1 min) by applying mild negative pressure (-50 mPa). This procedure allowed us to retain a tight seal during aspiration and to minimize RNA loss by keeping RNA molecules in solution within the pipette. When we broke contact, the recording pipette was pulled out from the recording chamber and then carefully rotated over an expelling 0.2 ml tube, where its content (0.5 μl) was ejected onto a 0.5 μl drop of lysis buffer pre-placed in the 0.2 ml tight-lock tube (TubeOne). The resultant sample (1 μl) was spun down (20 s) to the bottom of the conical tube at 24 °C, stored at -80 °C and later subjected to in-tube reverse transcription (RT).

#### Lysis, cDNA synthesis for library preparation and RNA-sequencing

Cell aspirates were dispensed into ∼0.5 μl lysis mix consisting of 0.15% Triton X-100, 1 U/μl TaKaRa RNase inhibitor, 2.5 mM dNTP, 17.5 mM DTT and oligo dT primer (2.5 uM). Samples were collected and stored at +80 °C until batch processing. Before reverse transcription, samples were thawed and lysed at 72 °C for 3 min, then cooled to 4 °C. Immediately following the lysis step, 2 μl RT mix (1× SuperScript II First-Strand Buffer; Life Technologies) supplemented with 15 mM MgCl2, 2.5 μM strand-switching primer “LNA”, 1.5 U/μl TaKaRa RNase inhibitor (Clontech), 1.45 M betaine and 21 U/μl Superscript II reverse transcriptase (Life Technologies)) were added and incubated at 42°C for 90 min followed by 72 °C for 10 min. Following reverse transcription, 8 μl PCR mix (1× KAPA HiFi 2× ready mix and 80 nM ISPCR primers) were added and PCR-amplified using thermal cycling as follows: 95 °C for 3 min (5 cycles), 98 °C for 20 s, 62 °C for 4 min, 72 °C for 6 min, (9 cycles) 98 °C for 20 s, 68 °C for 30 s, 72 °C for 6 min, (7 cycles) 98 °C for 30 s, 68 °C for 30 s, 72 °C for 7 min. Subsequently, PCR samples were cleaned using AMPure-XP beads (1:1 ratio; Beckman Coulter) and quantified by Qubit (Life Technologies) on an Agilent bioanalyzer. Tagmentation and subsequent library preparation were done using the Nextera-XT kit (Illumina). Paired-end (50bp) single-cell RNA-sequencing was done on an Illumina Hiseq2500 instrument, generating ∼1.5 million reads/cell. Alignment and locus mapping of the reads were done using HTseq and Bowtie 2.0.

#### Immunohistochemistry

For the anatomical analysis of the IG, guinea pig anti-*Vglut1*, rabbit anti-*Vglu2* and rabbit anti-*Vglut3* (1:1,000; Synaptic Systems, Cat. No. 135304; Cat. No. 135403; Cat. No. 135203, respectively), guinea pig anti-NPY (1:500; Abcam, Cat. No. ab10341), rabbit anti-glial fibrillary acidic protein (GFAP, 1:1,000; DAKO, Cat. No. GA52461-2), mouse anti-myelin basic protein (MBP, 1:200; R&D Systems, Cat. No. MAB42282), mouse anti-Iba1 (1:1,000, Millipore, Cat. No. MABN92), rabbit anti-*Scgn* (1:1,000, Atlas Antibodies, Cat. No. HPA006641), guinea pig anti-c-Fos (1,1:000, Synaptic Systems; Cat No. 226 005) and rabbit anti-*Brn*-1 (1:100, Santa Cruz, Cat. No. sc-6028) primary antibodies as well as *Solanum tuberosum* lectin (20 μg/ml; Vector Labs.) were employed according to standard protocols (2). Hoechst 33,342 (Sigma) was used as nuclear counterstain.

#### Tissue clearing of mouse samples, light-sheet microscopy

Freshly prepared brain slices of 500 μm were post-fixed in 4% PFA in phosphate buffer (PB, 0.1M, pH 7.8) at 4 °C overnight. Slices were repeatedly washed in PB and cleared using the CUBIC method (5) by first immersing in ‘CUBIC reagent 1’ (25% urea, 25% *N*,*N*,*N*′,*N*′-tetrakis(2-hydroxypropyl) ethylenediamine and 15% Triton X-100) for 2 d. *Scgn* immunohistochemistry started by blocking the specimens in 2% bovine serum albumin (BSA), 5% normal donkey serum (NDS) in 0.1M PB for 2h and then exposing those to rabbit anti-*Scgn* antibody (1:10,000) in 0.1% BSA, 1% NDS, 10% DMSO (to facilitate deep tissue penetration) on an orbital shaker at 22-24 °C for 3 days. This was followed by repeated rinses in 0.1M PB followed by exposure to a secondary antibody (Cy5-conjugated secondary antibody made in rabbit (1:200; Jackson) and at 22-24 °C for 2 days. Slices were then rewashed in PB and submerged in ‘CUBIC reagent 2’ (50 % sucrose, 25 % urea, 10% 2,20,20’-nitrilotriethanol and 0.1% Triton X-100) for further clearing. Cleared tissues from immunostainings or from *Fos*^CreER/T2^-based activity mapping were imaged on a laser-scanning microscope (Zeiss LSM880) or a Lightsheet.Z1 microscope (Zeiss).

Whole brains of E17.5 embryos were cleared using a modified CUBIC protocol (6). Brain samples were initially fixed in 4% PFA for 24h, and incubated in ‘CUBIC reagent 1’ under continuous agitation at 37 °C for 3 days. Subsequently, the samples were washed 3x 30 min each in 0.1M PB prior to immersion in ‘CUBIC reagent 2’ in which they were left at 22-24 °C for 3 days before imaging. All samples were imaged in ‘CUBIC reagent 2’ with a measured refractory index of 1.45. Orhtogonal images were acquired on a Lightsheet Z.1 (Zeiss) microscope using a 5× detection objective, 5× illumination optics and laser excitation at 488 nm. Each plane was illuminated from a single side, and images of the whole brain were obtained through 3 × 4 tile scanning. All images were captured at 0.7× zoom, with *z*-stack intervals set at 3.5 μm with an exposure time of 600 ms. 3D-rendered images were visualized with Arivis Vision4D for Zeiss (v. 2.12). Brightness and contrast of the 3D-rendered images were manually adjusted to aid visual clarity. The distribution of *ZsGreen1*^+^ cells was charted on coronal sections of both E17.5 (Prenatal mouse brain atlas (Schambra, Uta)) and adult brain templates (from The mouse brain atlas (Paxinos and Franklin) using overview brain Refs. Quantitative analysis was performed in Arivis.

#### Tract tracing

Twelve week-old rats were anesthetized (*i.m.*) with a mixture of ketamine (50 mg/kg) and xylazine (4 mg/kg) and placed in a Kopf stereotaxic instrument. The skull was exposed by a skin incision, and burr holes at corresponding coordinates were drilled to access the brain.

Tracers were injected stereotaxically using a 1.0 μl Hamilton syringe mounted on a microinjector (Kopf). Retrograde or anterograde tracers (40 nl each) were deposited by slow pressure injection during 5 min. The needle was retracted only after a 15 min resting interval to minimize leakage. Coordinates were verified in pilot experiments by methylene blue injections (Paxinos rat brain atlas). Alexa Fluor 594-conjugated choleratoxin B subunit (*retrograde* tracer, Molecular Probes; 1% dissolved in 0.05M phosphate-buffered saline) was injected into the olfactory bulb (AP: +7 mm, L: 1 mm, DV: -1.5 mm). Alexa Fluor-594 conjugated dextran (10 kDa, D594, Molecular Probes; 10% dissolved in distilled water) was infused into the fasciola cinerea (AP: -5.5 mm, L: -0.5 mm, DV: -2.5 mm). Four and 7 days after retrograde and anterograde tracing, respectively, rats were re-anesthetized with ketamine and xylazine as above and transcardially perfused first with 50 ml physiological saline solution followed by 250-300 ml of 4% PFA in 0.1M PB (pH 7.4).

The brains were post-fixed in 4% PFA in PB at 4 °C overnight, and transferred to 30% sucrose (diluted in PB, 4 °C, for 2 days) for cryoprotection. Brains were sectioned at 40 μm thickness on a Leica freezing microtome in the saggital plane. Free-floating sections were rinsed in PB (pH 7.4), blocked in 5% NDS (Jackson), and 0.3% Triton X-100 (Sigma) in PB for 2 h at 22-24 °C, then exposed (72 h at 4 °C) to a rabbit anti-*Scgn* antibody (1:10,000) in PB to which 0.1% NDS and 0.3% Triton X-100 had been added. After rinsing in PB, immunoreactivity was revealed by rabbit-specific Cy2 or Cy5-tagged secondary antibodies raised in donkey (1:500 (Jackson), 24 h at 4 °C). Glass-mounted sections were coverslipped with Surgipath Micromount (Leica).

#### Human adult samples

We have applied direct perfusion via the internal carotid and vertebral arteries, which facilitated the preservation of tissue integrity relative to alternative fixation methods. Human brains (*n* = 2, gender and age: female/83 years and male/79 years, with a clinical history of no neurodegenerative disease, ethical approval: TUKEB 84/2014, Hungary) were first perfused with physiological saline followed by a fixative containing 2% PFA and 0.1% glutaraldehyde in 0.1 M Tris-buffered saline (TBS, pH 7.4) 7h or 11h after death. The removal and subsequent preparation of human tissues were in accordance with relevant ethical guidelines of Semmelweis University (1998, Budapest, Hungary). Slices containing the IG were prepared from human tissue post-fixed in 2% PFA in TBS for 72, followed by immersion in cryoprotective 30% sucrose in 0.1M PB (pH 7.4) overnight. Coronal sections (50 μm) were cut on a cryostat microtome and processed for immunohistochemistry. Free-floating sections were rinsed in PB (pH 7.4) and pre-treated with 0.3% Triton-X 100 (in PB) for 1h at 22-24°C to enhance the penetration of antibodies. Non-specific immunoreactivity was suppressed by incubating our specimens in a cocktail of 5% NDS (Jackson), 10% BSA (Sigma) and 0.3% Triton X-100 (Sigma) in PB for 1h at 22-24 °C. Sections were exposed for up to 72h (at 4 °C) to the cocktail of primary antibodies (rabbit anti-*Scgn* (1:5,000); mouse anti-MAP2 (1:200; Sygma) and guinea pig anti-calretinin (1:1,000; Synaptic Systems) diluted in PB to which 0.1% NDS and 0.3% Triton X-100 had been added. After extensive rinsing in PB, immunoreactivities were revealed by chromogenic staining (as above) or by Cy2, 3 or 5-tagged secondary antibodies raised in donkey (1:200 [Jackson], 2h at 22-24 °C). Lipofuscin autofluorescence was quenched by applying Sudan Black-B (1%, dissolved in 70% ethanol (7)). Glass-mounted sections were coverslipped with Aquamount embedding medium (Dako). Sections were inspected and images acquired on a 710LSM confocal laser-scanning microscope (Zeiss) at 10x or 40x primary magnification, and pinhole settings limiting signal detection to 0.5-0.7 μm. Emission spectra for each dye were limited as follows: Cy2/505-530 nm, Cy3/560-610 nm, and Cy5/650-720 nm.

#### Human fetal tissue

Fetal brain tissue was obtained from spontaneous or medically-induced abortions. Only cases without genetic disorders, head injury, or neurological diseases were included. These cases showed no chromosome aberrations or *post-mortem* autolysis. Neuropathological examination excluded major central nervous system malformations, severe hypoxic/ischemic encephalopathy, intraventricular hemorrhages, severe hydrocephalus, and meningitis or ventriculitis. A 236 days old fetal brain with no disease history was then selected from the Brain Bank of the Institute of Neurology, Medical University of Vienna, Austria. All procedures were compliant with the Declaration of Helsinki and followed institutional guidelines. The study was approved by the Ethical Committee of the Medical University of Vienna (No.104/2009). Three-μm-thick sections from formalin-fixed, paraffin-embedded tissue blocks containing the corpus callosum and the hippocampal formation were mounted on pre-coated glass slides (Star Frost). Shortly after deparaffinization and rehydration, the sections were pretreated in low-pH EnVision FLEX antigen retrieval solution at 98 °C for 20 min (PTLink; Dako) and subsequently incubated with rabbit anti-*Scgn* antibody (1:12,000). The DAKO EnVision detection kit, peroxidase/3,3-diaminobenzidine-tetrahydrochloride (DAB), and rabbit secondary antibody (K5007, ready-to-use; Dako) were used to visualize antibody binding. Sections were counterstained with hematoxylin, dehydrated in ascending concentrations of ethanol, cleared with xylene, and covered with Consil-Mount (Thermo Scientific). Representative images containing the area of interest were exported from scanned sections (NanoZoomer 2.0; Hamamatsu).

#### Behavioral experiments

Methamphetamine (kind gift of Dr. C. Pifl, Medical University of Vienna) was injected *i.p.* at a dose of 15 mg/kg in male wild-type (*n* = 3) and *Scgn*^-/-^ mice (*n* = 4) (8-12 weeks of age) 3 times with 90 min intervals. All mice were tested on the same day in one cohort. The spontaneous behavioral state of each mouse was scored by 2 independent observers 30 min before the first injection (baseline) and 60 min after each methamphetamine injection using the SHIRPA protocol (8, 9). Criteria were set as follows: *body position*: 0 = inactive, 1 = active 4 paws, 2 = excessively active 2 paws, 3 = lordosis; *spontaneous activity*: 0 = none, 1 = slow movement, 2 = moderate movement, 3 = rapid movement; *respiration rate*: 0 = irregular, 1 = slow, 2 = normal, 3 = hyperventilation; *tremor*: 0 = absence, 1 = presence; *piloerection*: 0 = absence, 1 = presence; *gait*: 0 = normal, 1 = fluid but normal, 2 = limited movement, 3 = incapacity; *tail elevation*: 0 = dragging, 1 = horizontally extended, 2 = elevated, 3 = horizontally bent; *limb grasping*: 0 = absence, 1 = present in forelimb, 2 = present in hindlimb, 3 = limb rigidity/less extended; *clonic convulsions*: number of clonic convulsions in 60 sec; *tongue stuck out*: 0 = absence, 1 = less/licking the wall on 2 paws, 2 = almost continuous. Sample videos were taken after the scoring of behavioral state at 60 min indicated as test two (T2).

#### Figures and statistics

Composite figures were assembled in CorelDraw X7 (Corel Corp.). Statistical analysis of data for electrophysiology and behavioral experiments was carried out in Matlab. We used the Mann-Whitney *U*-test to determine group differences whenever the Shapiro-Wilk test failed to verify normal distribution of the data; including methamphetamine-induced modifications in *Scgn*^-/-^ mice. Contrast analysis to identify IG-specific neuronal markers in *Patch-seq* experiments was also performed in Matlab. We used ‘reads per kilo base per million mapped reads’ (RPKM) data as Tasic et al. (10) to aid comparative assessment. We filtered the set of genes obtained from visual cortical neurons with zero or low expression (low-pass filtering) and compared these to highly-expressed genes in IG from our *Patch-seq* dataset (high-pass filtering). The open-source Allen brain atlas adult mouse *in situ* hybridization database was then used to manually filter putative marker genes.

## Legends to Supplementary Figures & Videos

**Figure S1:**
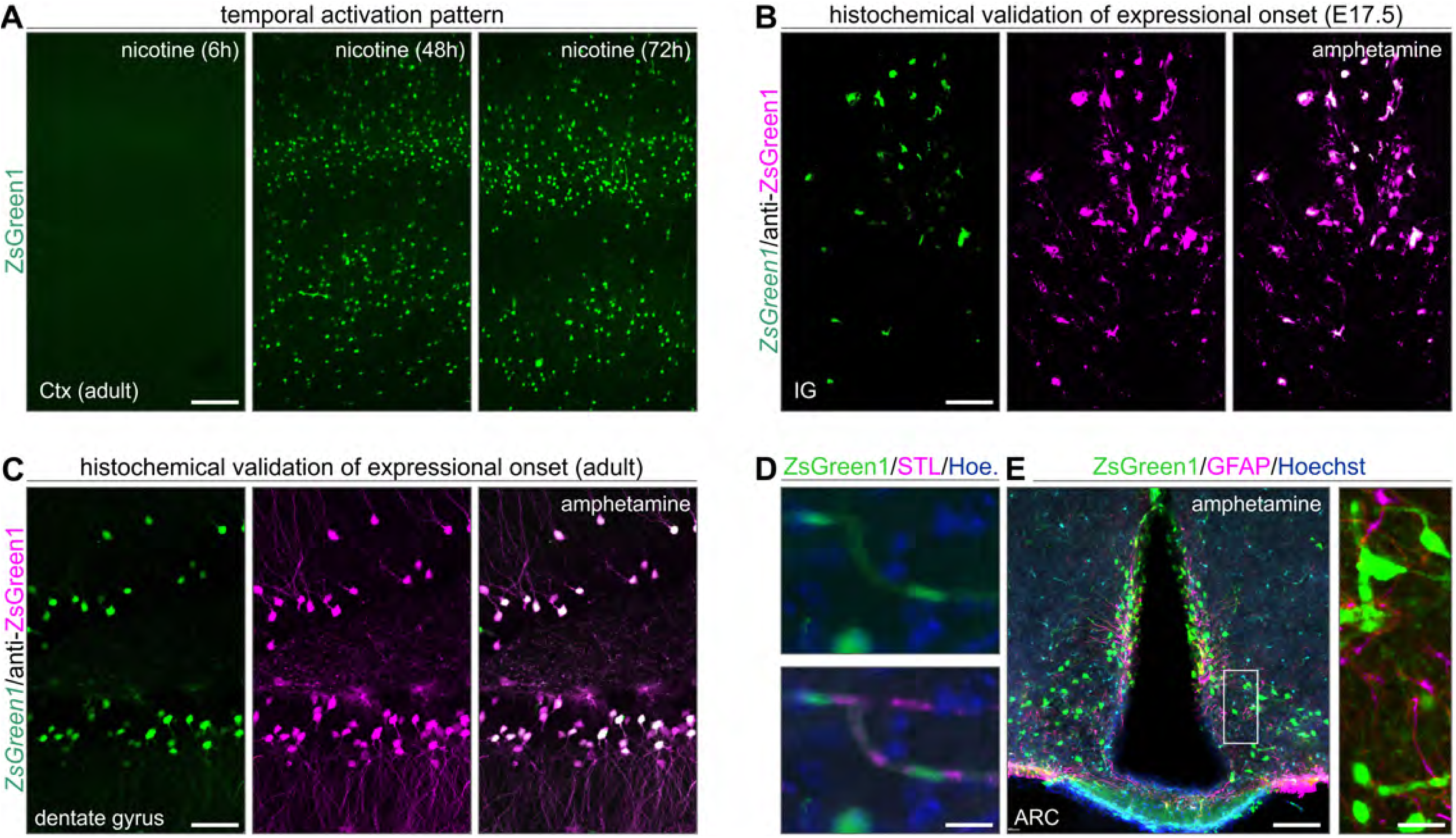
Methodological controls of the transgenerational ‘TRAP’ mouse model. (**A**) Considering that immediate early gene-CreER^T2^-dependent models to detect transient neuronal activity rely on the coincidence of tamoxifen-induced Cre activity and the experimental manipulation, it is imperative to control if the temporal setting of the experiments allows for successful recombination. Here, nicotine is shown to induce recombination in the neocortex, with abundant *ZsGreen1* expression by 48h and 72h. (**B,C**) Next, an anti-ZsGreen1 antibody with indirect fluorescence detection was used in both fetal (B) and adult (C) brains to show a virtually complete overlap with genetically-encoded ZsGreen1 fluorescence. In both cases, amphetamine was used to trigger cellular activity. (**D**) Besides neurons, ZsGreen1 signal was also seen in endothelial cells, which were labeled by *Solanum tuberosum* lectin (STL). Nuclei were counterstained with Hoechst 33,342. (**E**) Amphetamine-induced neuronal activation in the arcuate nucleus (ARC). Note that *ZsGreen1* fills release terminals at the median eminence. Inset shows the relative abundance of *ZsGreen1* in neurons but not astroglia (labeled for glial fibrillary acidic protein; GFAP). *Abbreviations*: Ctx, cortex; IG, indusium griseum; *Scale bars* = 200 μm (A), 50 μm (C), 30 μm (B).

**Figure S2:**
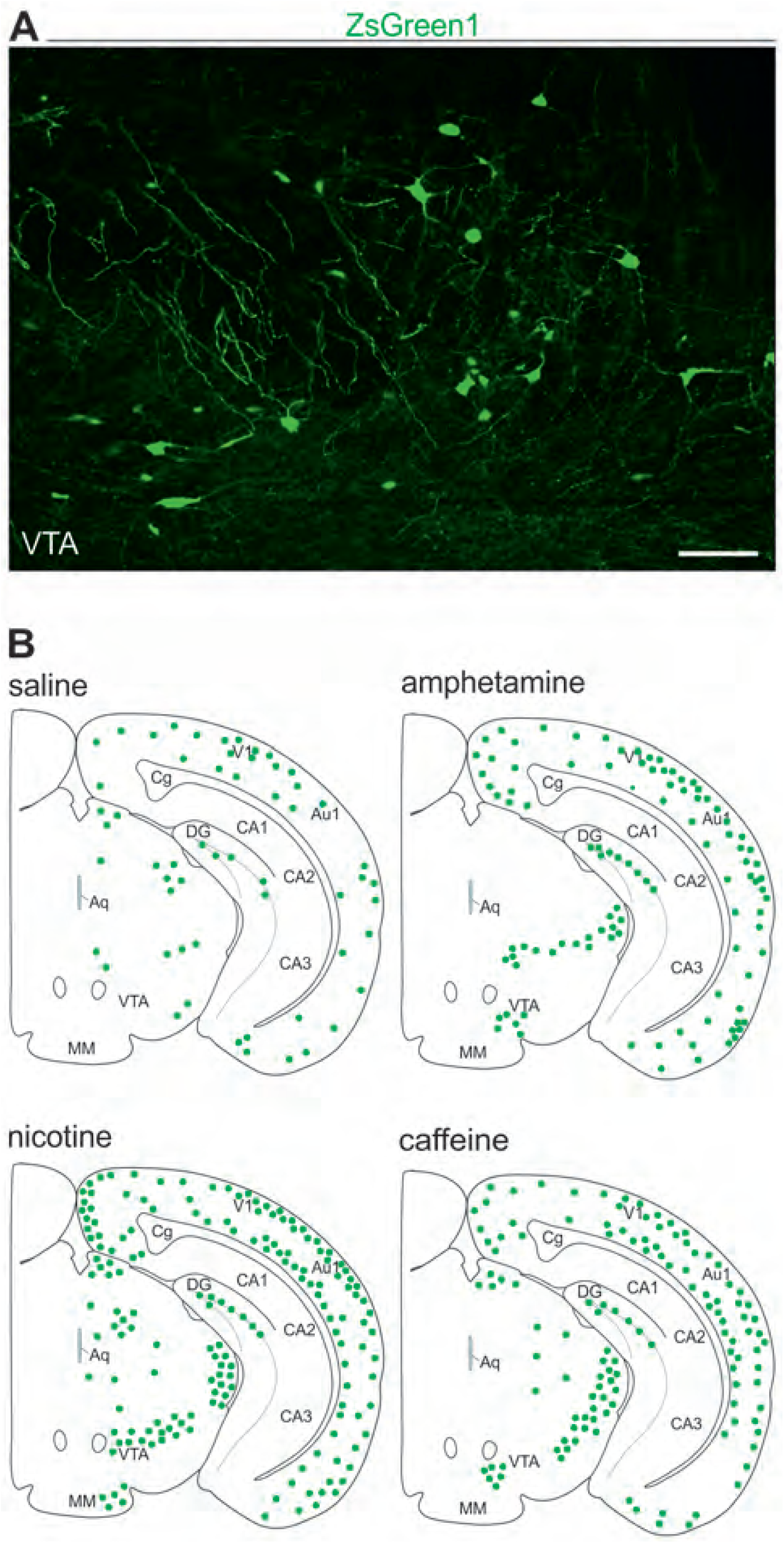
Neuronal activation in the ventral tegmental area upon psychostimulant exposure. (**A**) In adult *Fos*-CreER^T2^::*ZsGreen1*^stop-floxed^ mice, all three psychostimulants used activated neuronal contingents in the vental tegmental area (VTA). Here, the effects of amphetamine are shown. (**B**) Distribution of *ZsGreen1*^+^ neurons after amphetamine, nicotine and caffeine treatment at the coronal plain spanning the VTA. These observations served as positive control for the drug effects. *Abbreviations*: Aq, aqueduct; Au1, auditory cortex; CA1-3, Cornu Ammonis 1-3 subfields of the hippocampus; DG, dentate gyrus; MM, medial mammillary nucleus; VTA, ventral tegmental area; V1, visual cortex. *Scale bar* = 75 μm.

**Figure S3:**
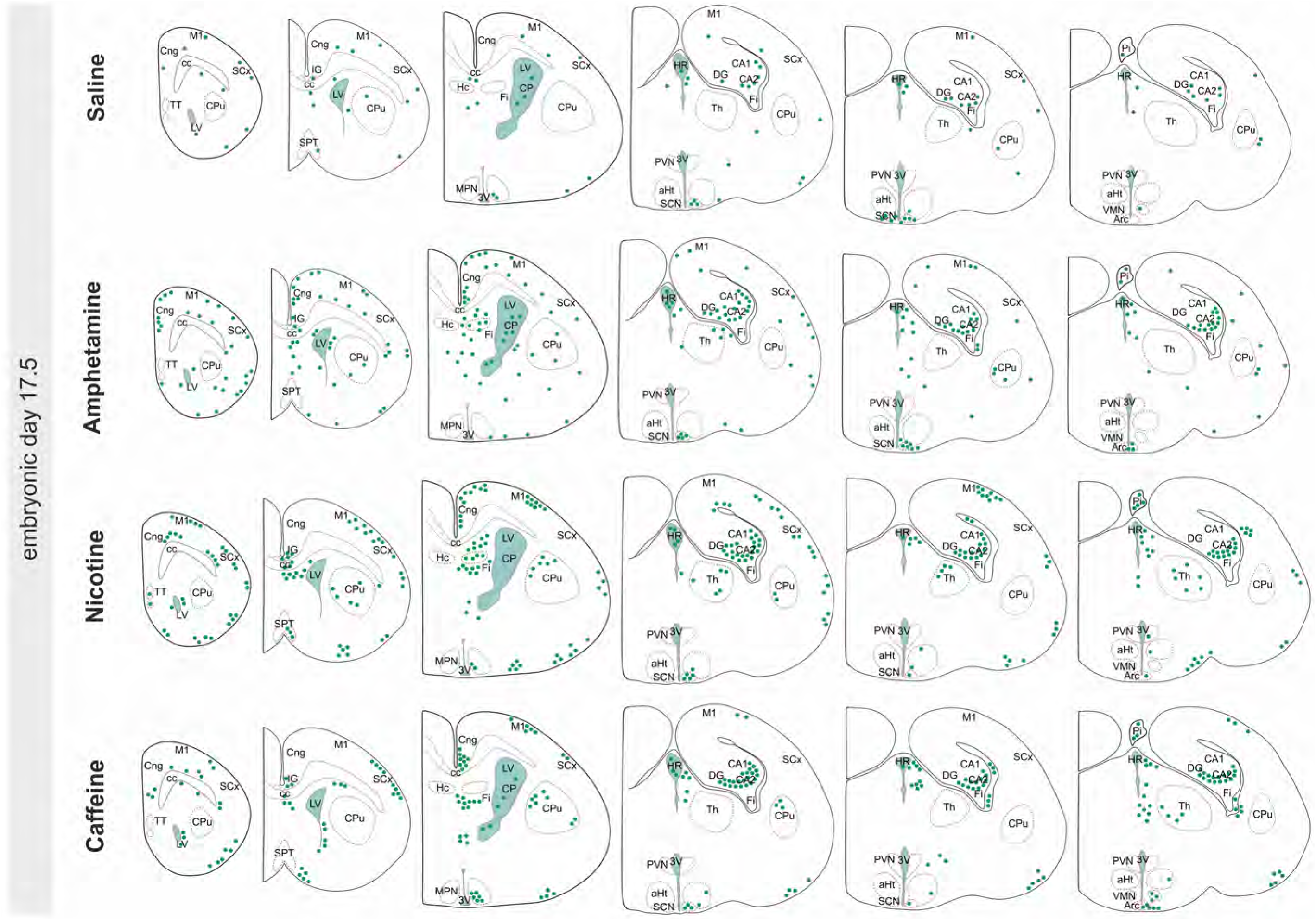
Cell activation maps for fetal brains. *Fos*-CreER^T2^::*ZsGreen1*^stop-floxed^ mice were treated with the psychostimulants as described (Fig. 1A). After 72h survival, serial brain sections were analyzed to localize *ZsGreen1*^+^ somata (without histochemical signal amplification). Data are from *n* ≥ 4 fetal brains/treatment, which were mapped by at least two independent researchers blinded to the case conditions. *Abbreviations*: aHt, anterior hypothalamus; ARC, arcuate nucleus; cc, corpus callosum; CA1,2, Cornu Ammonis 1,2 subfields of the hippocampus; Cng, cingulate cortex; CP, choroid plexus; CPu, striatum; DG, dentate gyrus; Fi, fimbria hippocampi; Hc, hippocampus; HR, habenular recess; IG, indusium griseum; LV, lateral ventricle; M1, motor cortex; MPN; median preoptic nucleus; Pi, pineal gland; PVN, parsventricular nucleus of the hypothalamus; SCN, suprachiasmatic nucleus; SCx, somatosensory cortex; SPT; septum; Th, thalamus; TT, tenia tecta; VMN, ventromedial hypothalamic nucleus; 3V, 3^rd^ ventricle.

**Figure S4:**
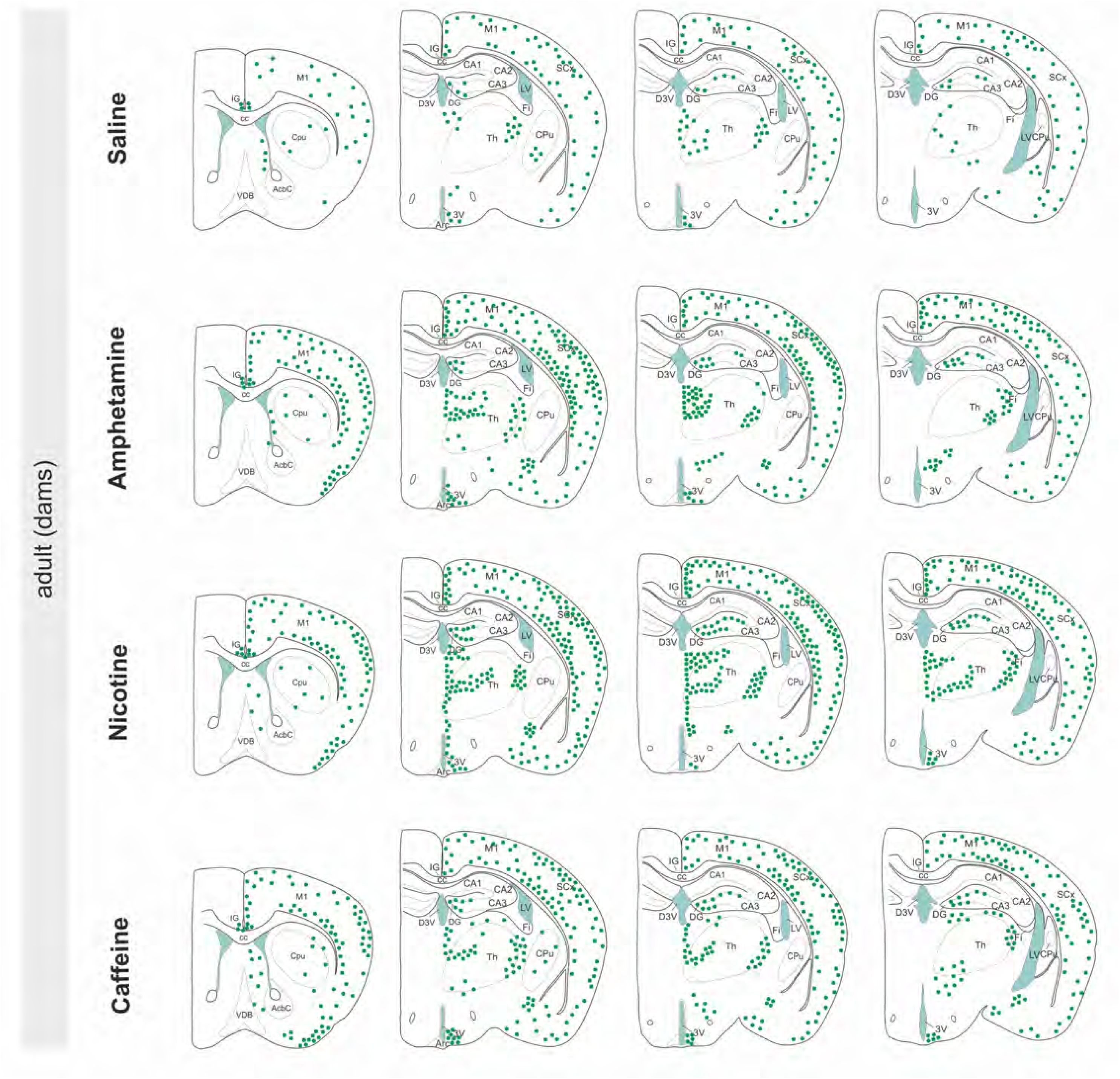
Cell activation maps for (adult) dams. *Fos*-CreER^T2^::*ZsGreen1*^stop-floxed^ mice were treated with the psychostimulants as described (Fig. 1A). After 72h survival, serial brain sections were analyzed to localize *ZsGreen1*^+^ somata (without histochemical signal amplification). Data are from *n* ≥ 3 animals/treatment, which were mapped by at least two independent researchers blinded to the case conditions. *Abbreviations*: AcbC, core of the nucleus accumbens; ARC, arcuate nucleus; CA1-3, Cornu Ammonis 1-3 subfields of the hippocampus; cc, corpus callosum; CPu, striatum; D3V, dorsal part of the 3^rd^ ventricle; DG, dentate gyrus; Fi, fimbria hippocampi; IG, indusium griseum; LV, lateral ventricle; M1, motor cortex; SCx, somatosensory cortex; Th, thalamus; VDB, vertical diagonal band of Broca; 3V, 3^rd^ ventricle.

**Figure S5:**
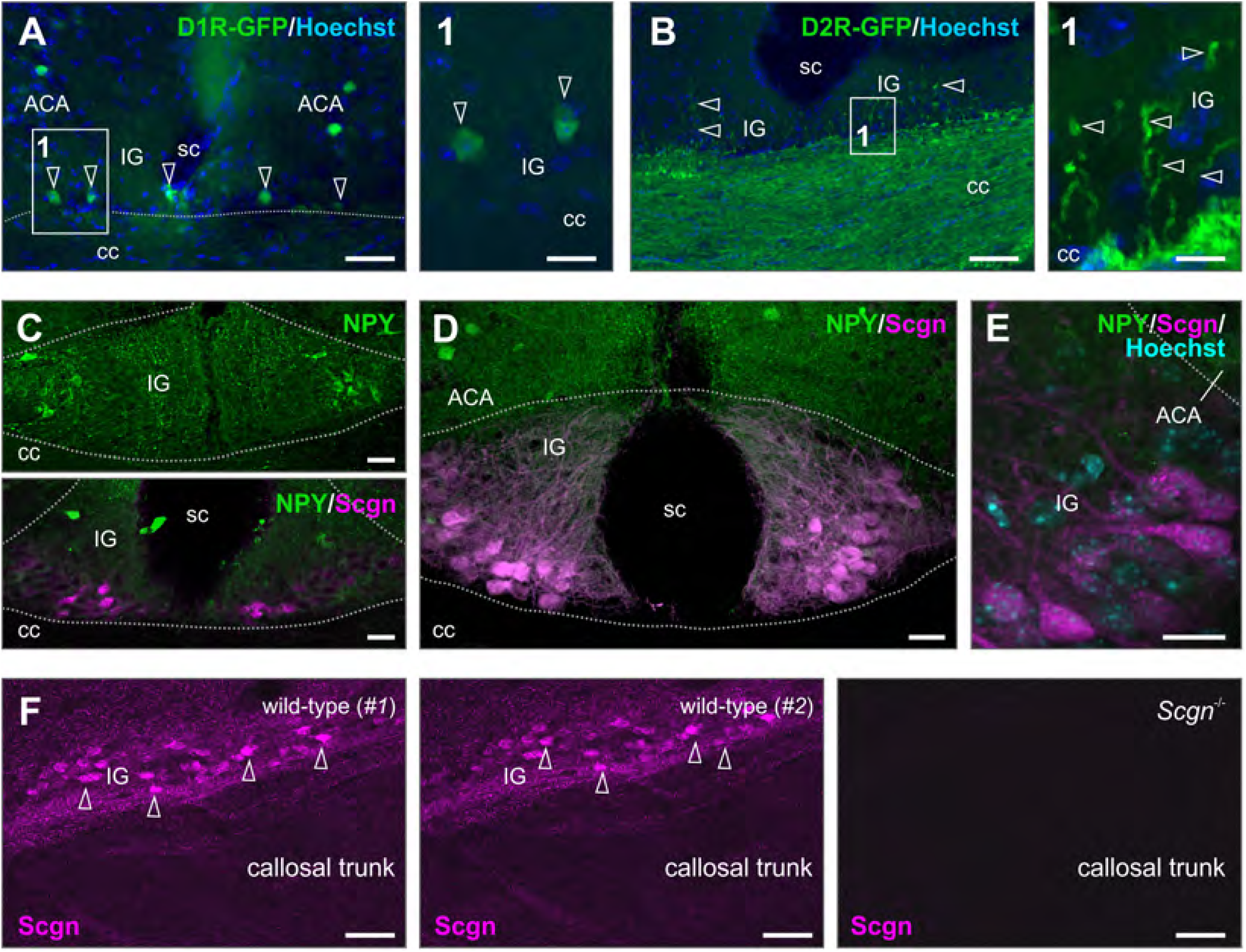
Neuropeptide Y (NPY) immunoreactivity during IG development. (**A,B**) Dopamine receptor 1 (D1R) and 2 (D2R) promoter controlled GFP expression in *D1R*-GFP (**A**) and *D2R*-GFP (**B**) mouse lines labels IG neurons and callosal afferents (*open arrowheads*), respectively. Enumerated rectangles denote the location of insets. Sections were counterstained with Hoechst 33,342. (**C**) NPY immunoreactivity in the putative IG on P5 shows transient neuropeptide expression in lateral cell columns (*top*). *Scgn* is not expressed in the IG at this time (*not shown*). By P12, the IG is NPY^-^ with a few cells gaining *Scgn* immunoreactivity (*bottom*). (**D**) On P25, the IG remains essentially negative for NPY (close-up in **E**) with *Scgn*^+^ neurons forming dense clusters on its ventral surface adjacent to the corpus callosum. (**F**) *Scgn* immunolabeling of wild-type and *Scgn*^-/-^ IG supports the specificity of the novel antibody used here. *Abbreviations*: ACA, anterior cingulate area; cc, corpus callosum; IG, indusium griseum; P, postnatal day; sc sulcus centralis. *Scale bars* = 75 μm (A,B,F), 30 μm (C,D), 20 μm (A1,E), 12 μm (B1).

**Figure S6:**
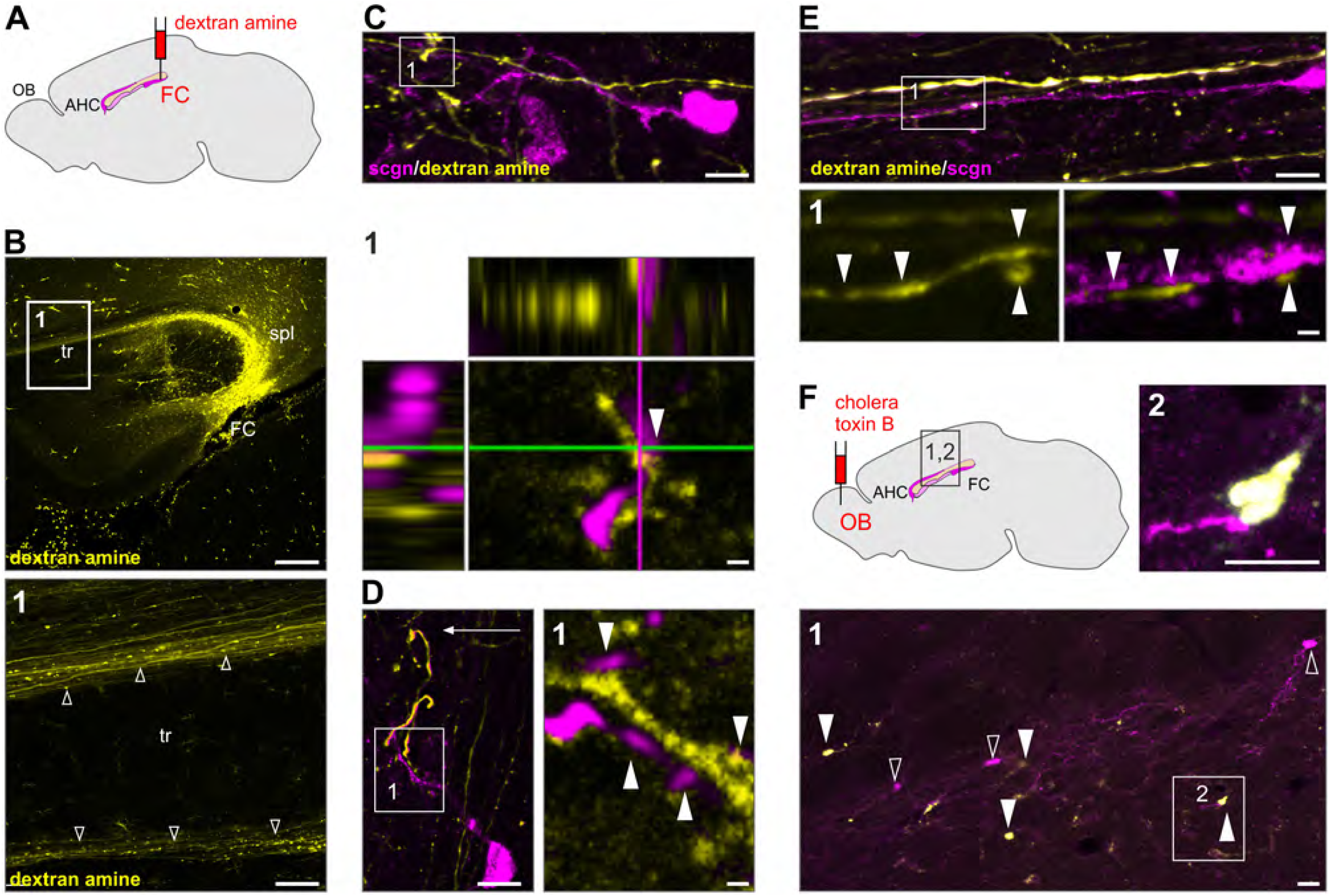
Circuit organization of *Scgn*^+^ IG neurons. (**A**) Dextran amine was used as anterograde tracer and injected into the fasciola cinerea (FC) of rats. (**B**) As a result, fluorescently-labelled axons ran along the dorsal and ventral surfaces of the corpus callosum (*open arrowheads*). (**C-D**) Bouton-like fluorescently-labelled specializations were seen in close apposition to *Scgn*^+^ dendrites in the IG (arrowheads in C’1’, D). The lack of appreciable physical separation between pre- (dextrane amine^+^) and postsynaptic (*Scgn*^+^) neuronal compartment was verified by high-resolution orthogonal imaging (**C’1’**). (**E**) *Scgn*^+^ dendrites were oriented parallel with anterogradely-traced axons and were contacted by *en passant* boutons (*arrowheads*, ‘1’). (**F**) Injection of cholera toxin B, a retrograde tracer, into the olfactory bulb (OB) labeled *Scgn*^+^ neurons in the IG (‘1’,’2’). *Abbreviations:* AHC, anterior hippocampal continuation; Scgn, secretagogin. *Scale bars* = 200 μm (B,B’1’), 50 μm (F’1’), 20 μm (E), 10 μm (C,D,F’2’), 2 μm (C’1’, D’1’,E’1’).

**Figure S7:**
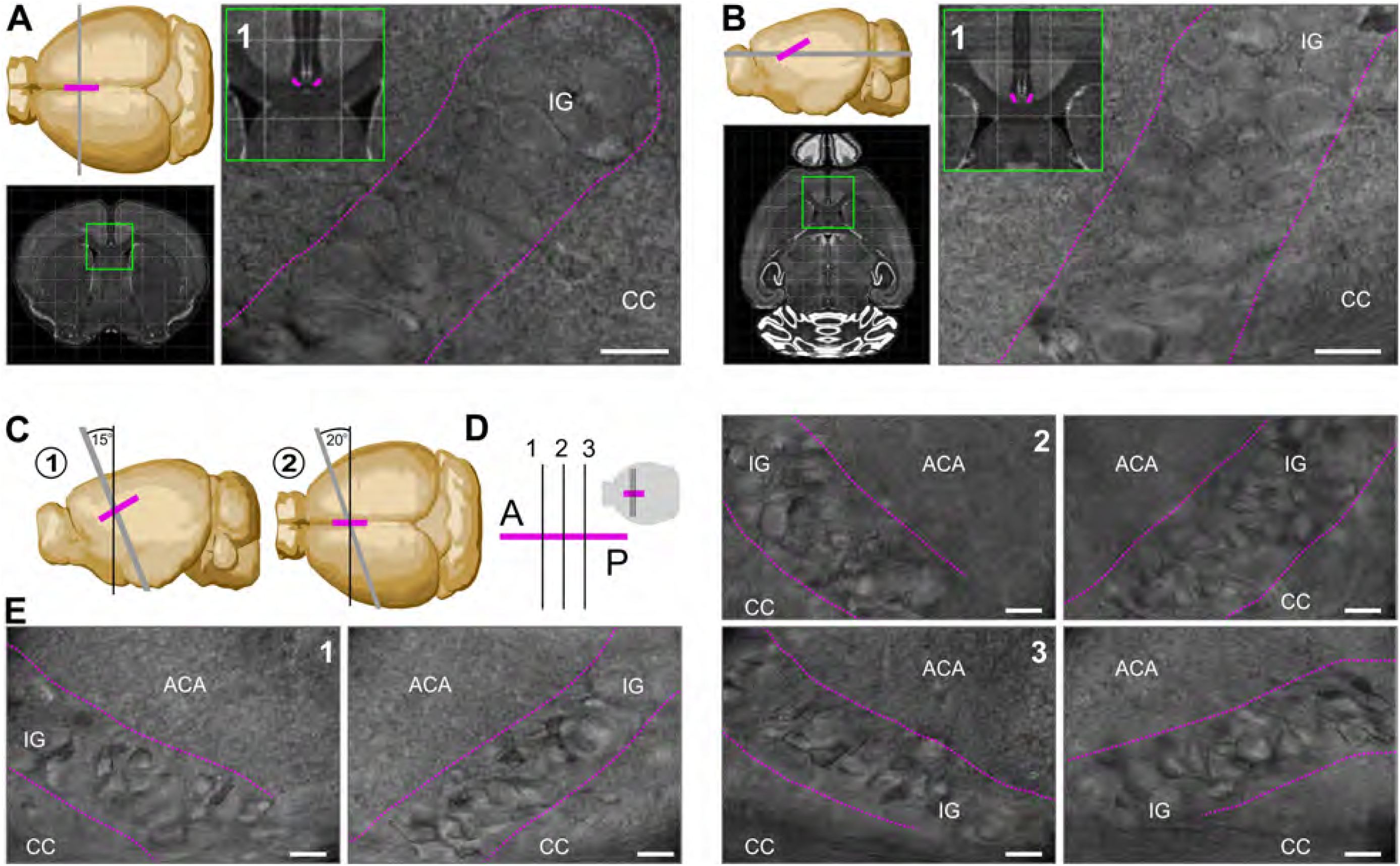
Cutting angle for improved neuronal survival in IG during slice preparation. (**A**) Coronal *ex vivo* brain slice preparation. Schema illustrates the coronal cutting plane (*top left*) and ensuing coronal section (*bottom left*) with the IG in magenta. (**’1’**) Differential interference microscopy (DIC) image of indusium griseum (IG) neurons in a coronal brain slice (dashed magenta line shows the boundary of this cell cluster) from the anatomical region indicated (*inset*; modified from ‘Brain Explorer’ of the Allen Institute). (**B**) Horizontal *ex vivo* brain slice preparation. The horizontal cutting plane (*top left*) and ensuing horizontal section (*bottom left*) are shown with magenta color indicating the location of the IG. (**‘1’**) DIC image of IG neurons in a horizontal brain slice (dashed magenta line shows their physical boundary) from the anatomical region indicated (*insert*). (**C**) Para-coronal brain slice preparation. Cartoon illustrates the two steps of para-coronal brain sectioning (‘1’ and ‘2’). (**D**) Schema with the position (albeit without orientation) of para-coronal cutting planes shown in E. (**E**) DIC image of para-coronal brain slices with IG (*dashed magenta line*) in slices from the most anterior (‘1’) towards posterior (‘3’) coordinates. *Scale bars* = 10 μm.

**Figure S8:**
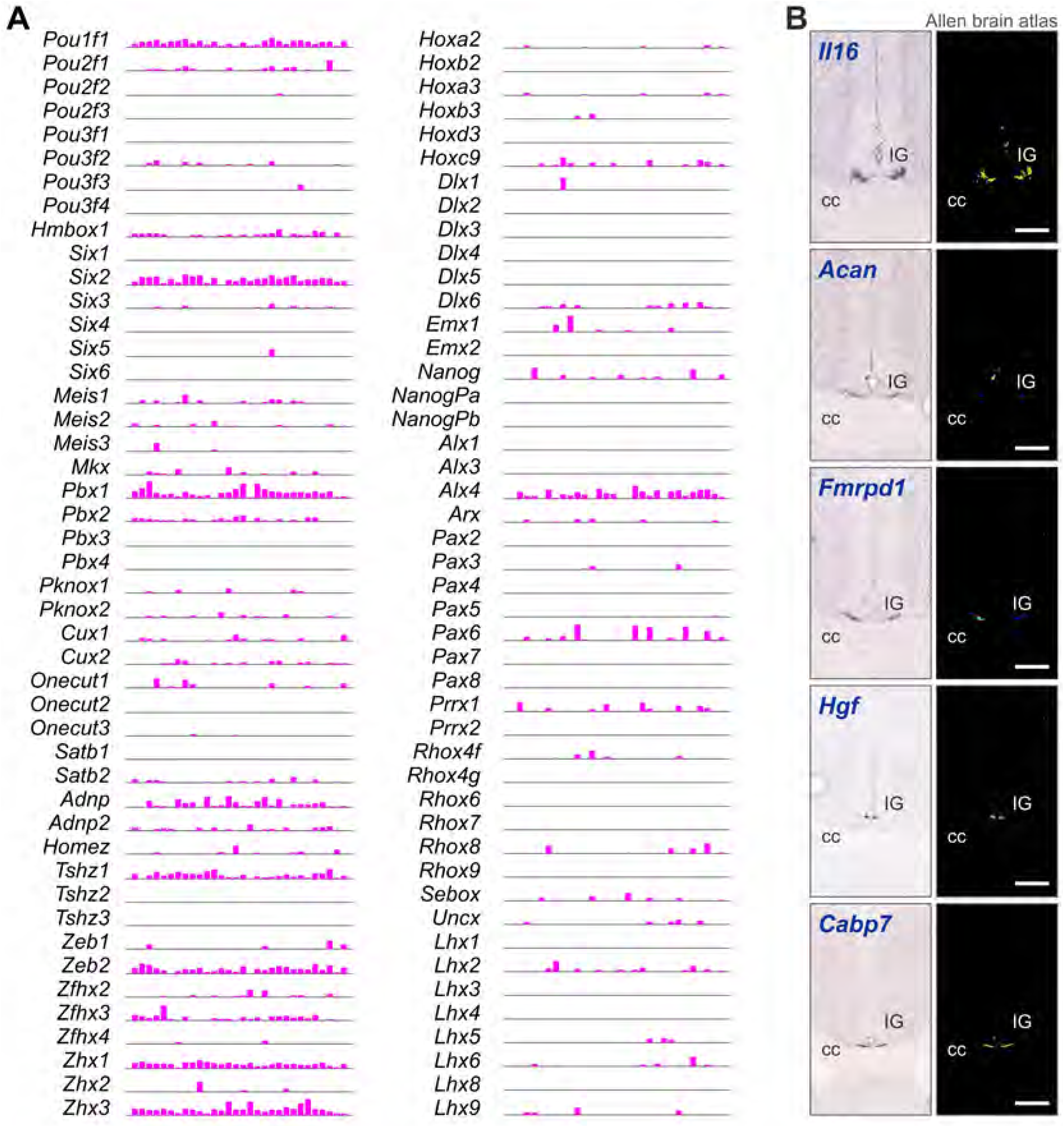
Transcription factor families and IG-specific markers. (**A**) Transcription factors in IG neurons probed by *Patch-seq*. Transcription factor families were plotted if at least one of the family members was detected in >15% of the IG neurons analyzed. (**B**) IG-specific gene expression deduced from our *Patch-seq* data with the Tasic *et al*. (2016) cortical single-cell RNA-seq dataset used as reference. IG-specific enrichment was verified by surveying open-source *in situ* hybridization results from the Allen Institute (www.brain-map.org).

**Figure S9:**
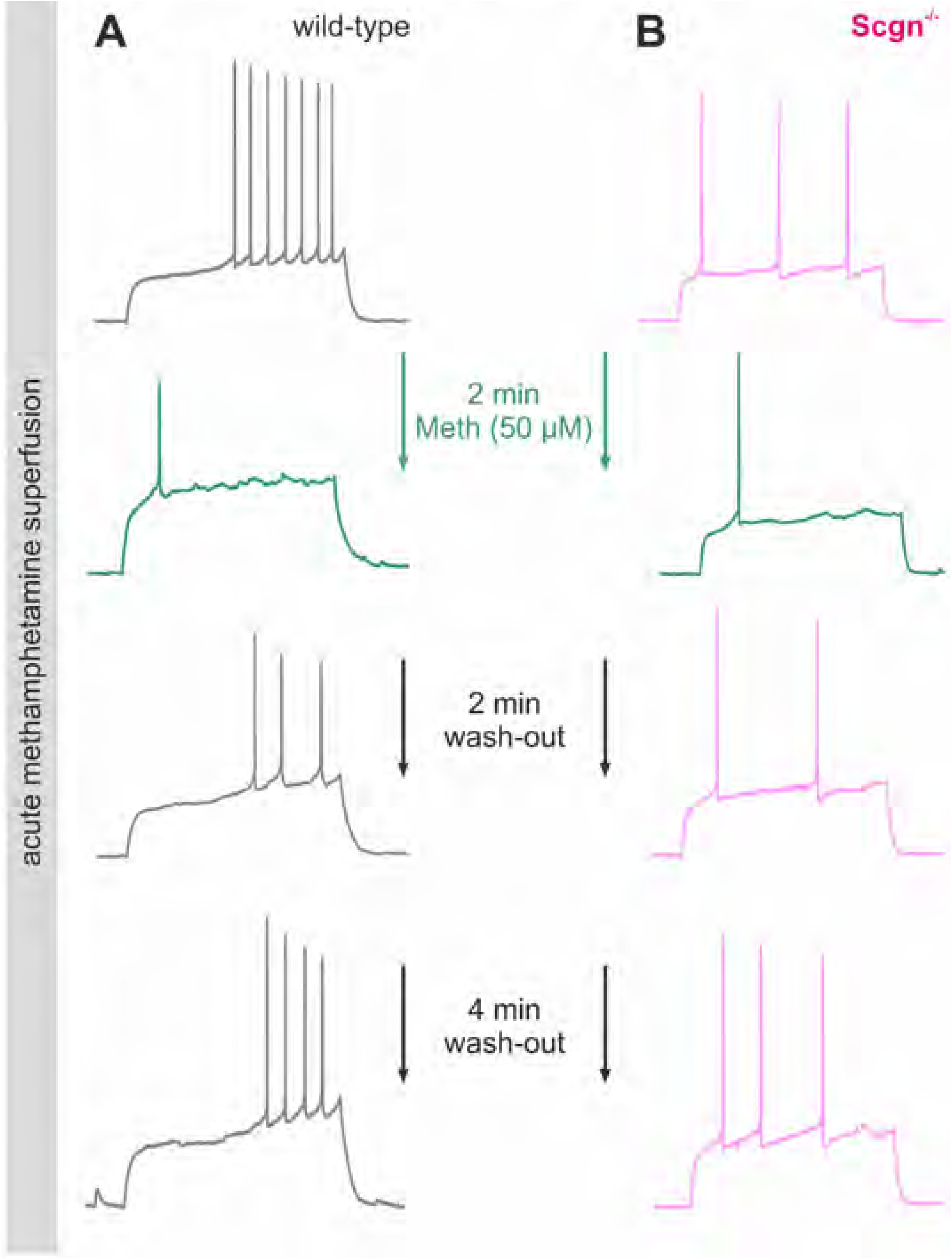
Acute methamphetamine pharmacology on wild-type and *Scgn*^-/-^ IG neurons. Examples of neuronal firing upon exposure to 50 uM methamphetamine (*Meth*) superfused in the slice recording chamber. (**A)** *Meth* application on a wild-type IG neuron resulted in reduced AP formation within 2 min of superfusion. The reversible nature of *Meth* effect was verified by wash-out after 2 min. At 4 min, the firing pattern of IG neuron was comparable to that during the baseline. (**B)** *Meth* application onto *Scgn*^-/-^ IG neurons resulted in an effect similar to that seen in wild-type neurons along with a comparable wash-out kinetics suggesting that *Scgn* does not impede *Meth* responsiveness acutely.

**Figure S10:**
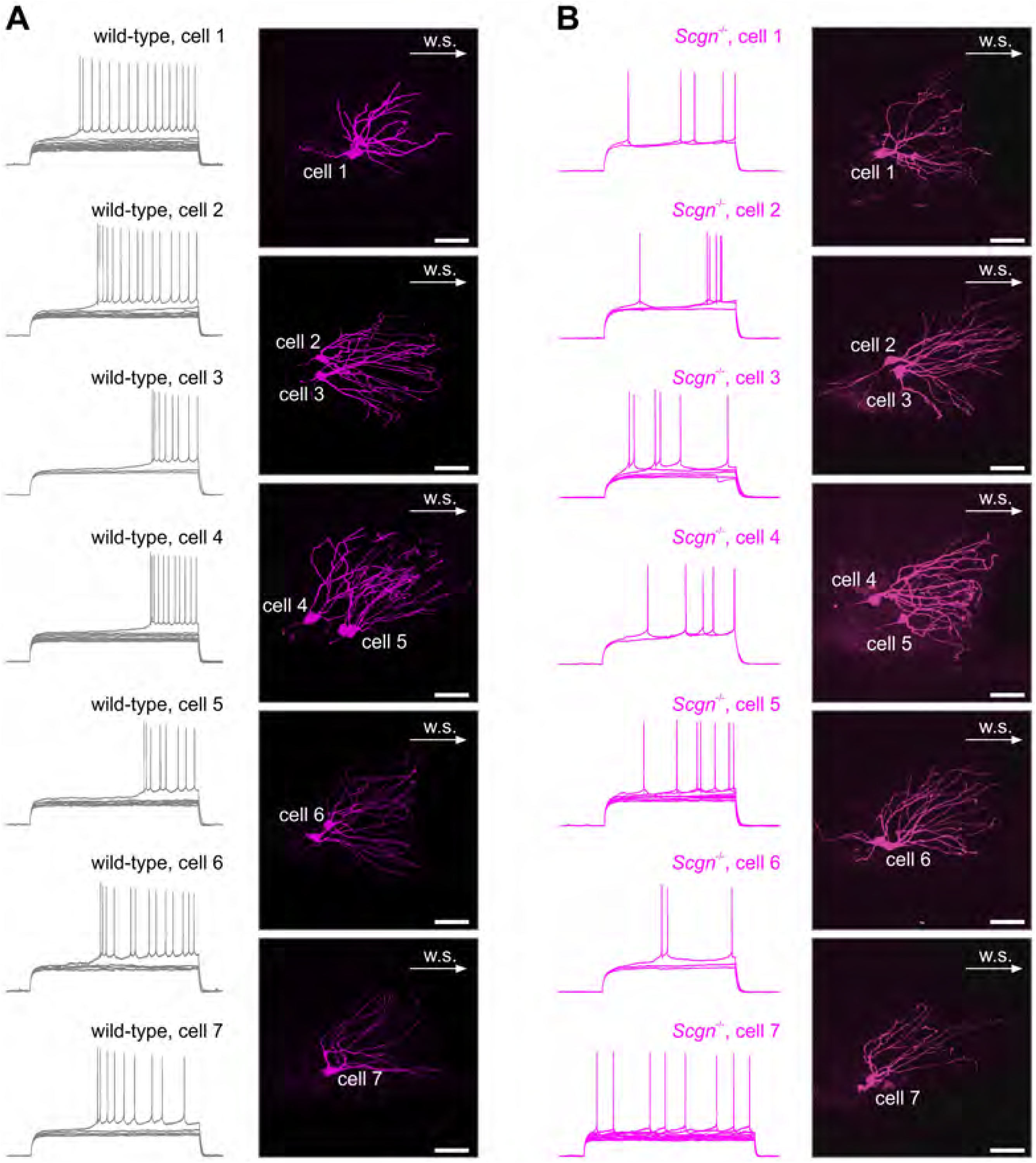
Examples of rheobasic firing of wild-type *vs*. *Scgn*^-/-^ IG neurons. (**A**) Wild-type IG neurons displayed a delayed rheobasic activation initiated by a pre-potential (that is, a pronounced depolarization to supra-threshold membrane potential resulting in a train of APs; *left*). Corresponding neuronal morphology is shown on the right. (**B**) In contrast, a pre-potential was not appreciable in *Scgn*^-/-^ IG neurons and consequently their discharge with lower rheobasic frequencies. *Scgn*^-/-^ IG neuronal morphology is shown to the right; *n* = 7 neurons were selected from both groups; current steps = 2 s; *scale bars* = 20 um, arrows indicate picture orientation and point towards the midline of the brain.

**Figure S11:**
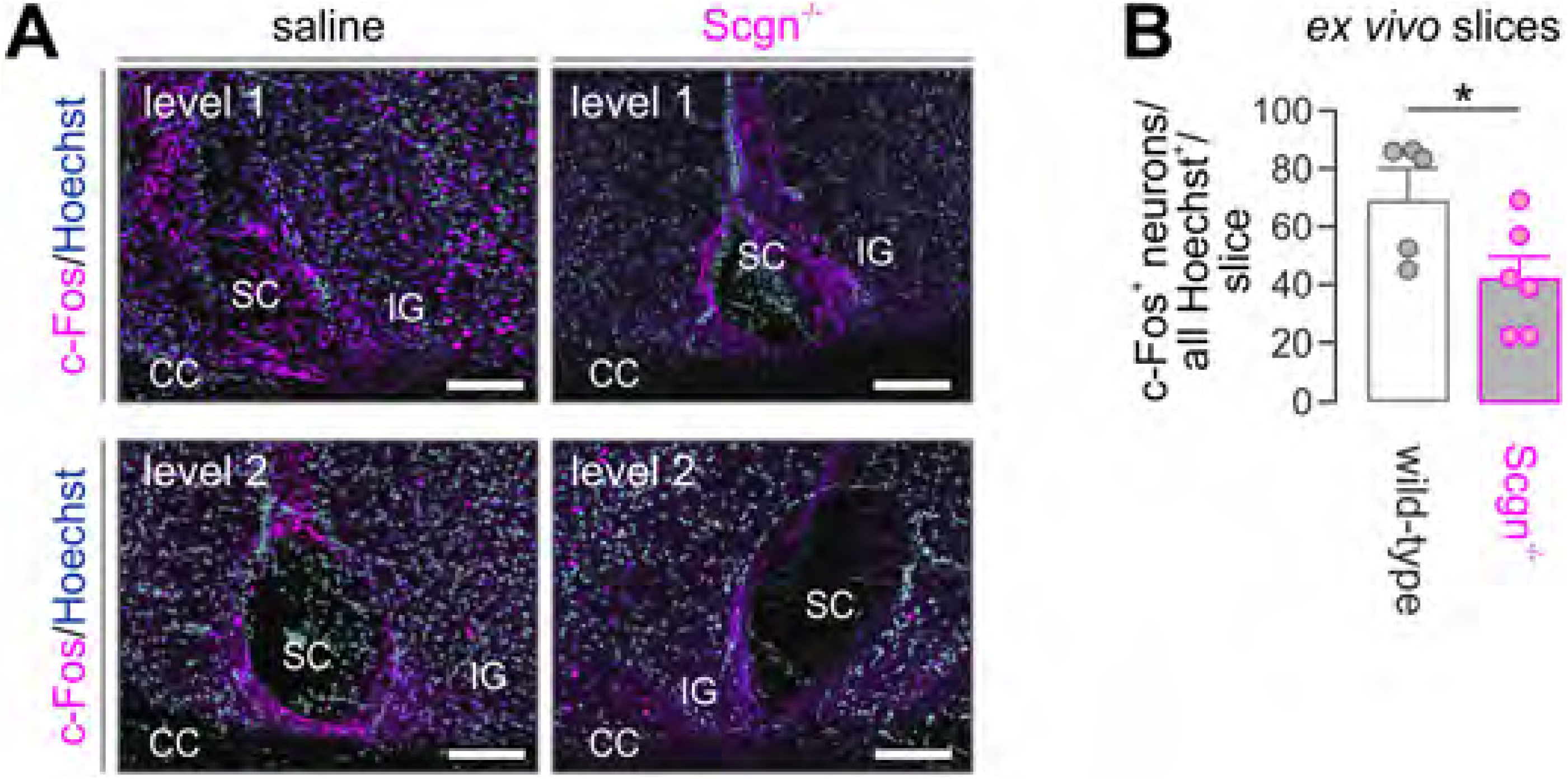
C-Fos activation in the postnatal IG challenged with methamphetamine. (**A**) Injection (*i.p*.) of saline (*left*) or methamphetamine (*Meth*, *right*) resulted in a low *vs*. high IG neuronal activation respectively, histochemically visualized by c-Fos immunoreactivity. (**B**) Quantification of c-Fos expression as a ratio of c-Fos^+^/all IG nuclei (labeled by Hoechst staining). Data were expressed as means ± s.e.m., *n* = 3 animals/group; \**p* < 0.05 (Student’s *t*-test).

**Videos S1-4:** Lightsheet microscopy of fetal brains after ZsGreen activation.

**Videos S5,6:** Methamphetamine-induced behaviors of wild-type and *Scgn*^-/-^ mice.

Video files are available for viewing/download at: https://www.dropbox.com/login?src=logout Login ID/ Password: Fuziketal2019

